# Asymmetric roles for M2-M3 linkers of specific GABA_A_ receptor subunits in the intrinsic closed-open equilibrium and its modulation by the benzodiazepine diazepam

**DOI:** 10.1101/2023.10.30.564751

**Authors:** Joseph W. Nors, Zachary Endres, Marcel P. Goldschen-Ohm

## Abstract

GABA_A_ receptors (GABA_A_Rs) are neurotransmitter-gated ion channels critical for inhibitory synaptic transmission as well as the molecular target for benzodiazepines (BZDs), one of the most widely prescribed class of psychotropic drugs today. Despite structural insight into the conformations underlying functional channel states, the detailed molecular interactions involved in conformational transitions and the physical basis for their modulation by BZDs are not fully understood. We previously identified that alanine substitution at the central residue in the α1 subunit M2-M3 linker (V279A) enhances the efficiency of linkage between the BZD site and the pore gate. Here, we expand on this work by investigating the effect of alanine substitutions at the analogous positions in the M2-M3 linkers of β2 (I275A) and γ2 (V290A) subunits, which together with α1 comprise typical heteromeric α1β2γ2 synaptic GABA_A_Rs. We find that these mutations confer subunit-specific effects on the intrinsic pore closed-open equilibrium and its modulation by the BZD diazepam (DZ). The mutations α1(V279A) or γ2(V290A) bias the channel toward a closed conformation, whereas β2(I275A) biases the channel toward an open conformation to the extent that the channel becomes leaky and opens spontaneously in the absence of agonist (e.g., GABA). In contrast, only α1(V279A) enhances the efficiency of DZ-to-pore linkage, whereas mutations in the other two subunits have no effect. These observations show that the central residue in the M2-M3 linkers of distinct subunits in synaptic α1β2γ2 GABA_A_Rs contribute asymmetrically to the intrinsic closed-open equilibrium and its modulation by DZ.

## Introduction

GABA_A_Rs are the primary neurotransmitter-gated ion channels mediating inhibitory synaptic signaling throughout the central nervous system (Smart & Stephenson, 2019). They belong to the superfamily of cys-loop pentameric ligand-gated ion channels (pLGICs) including glycine, nicotinic acetylcholine, and serotonin receptors (Pless & Sivilotti, 2019). Genetic mutations conferring GABA_A_R dysfunction are associated with human disorders including epilepsy, autism spectrum disorder, intellectual disability, schizophrenia, and neurodevelopment disorders such as fragile X syndrome (Braat & Kooy, 2015; Gao et al., 2018; Ghit et al., 2021; Hernandez & Macdonald, 2019; Mahdavi et al., 2018; Ramamoorthi & Lin, 2011; Solís-Añez et al., 2007). GABA_A_Rs are also the molecular target for numerous anxiolytic, analgesic, and sedative compounds. One of the most widely prescribed classes of psychotropic drugs are benzodiazepines (BZDs) (Agarwal & Landon, 2019), whose modulation of GABA_A_R activity is used to treat neurological conditions including anxiety, insomnia, muscle spasms, pain, and epilepsy (Möhler et al., 2002). Many functional studies together with recent cryogenic electron microscopy (cryo-EM) structural models of heteromeric synaptic GABA_A_Rs have begun to paint a picture of the conformational landscape involved in channel gating. However, a comprehensive view of the molecular details that govern the energetics of conformational transitions remains incomplete. Furthermore, the mechanism for allosteric modulation by drugs such as BZDs are only poorly understood (Goldschen-Ohm, 2022).

Typical synaptic GABA_A_Rs are hetero-pentamers comprised of α1, β1-3, and γ2 subunits **(Figure 1)** (Olsen & Sieghart, 2008; Sieghart & Sperk, 2002). The M2-M3 linker at the interface between the extracellular domain (ECD) and the transmembrane domain (TMD) has been identified as an important region for transducing the chemical energy from agonist (e.g., GABA) binding in the ECD to gating of the channel pore in the TMD **(Figure 1)** (Jha et al., 2007; Kaczor et al., 2022; Kash et al., 2003; O’Shea & Harrison, 2000; Sigel et al., 1999). Structurally, the M2-M3 linker moves radially outward along with the top of the pore-lining M2 helices during channel opening (Kim et al., 2020; Masiulis et al., 2019; Nemecz et al., 2016). However, the role of the M2-M3 linker in allosteric modulation by BZDs or the intrinsic energetics of the pore in the absence of agonist are less studied.

**Figure 1.**
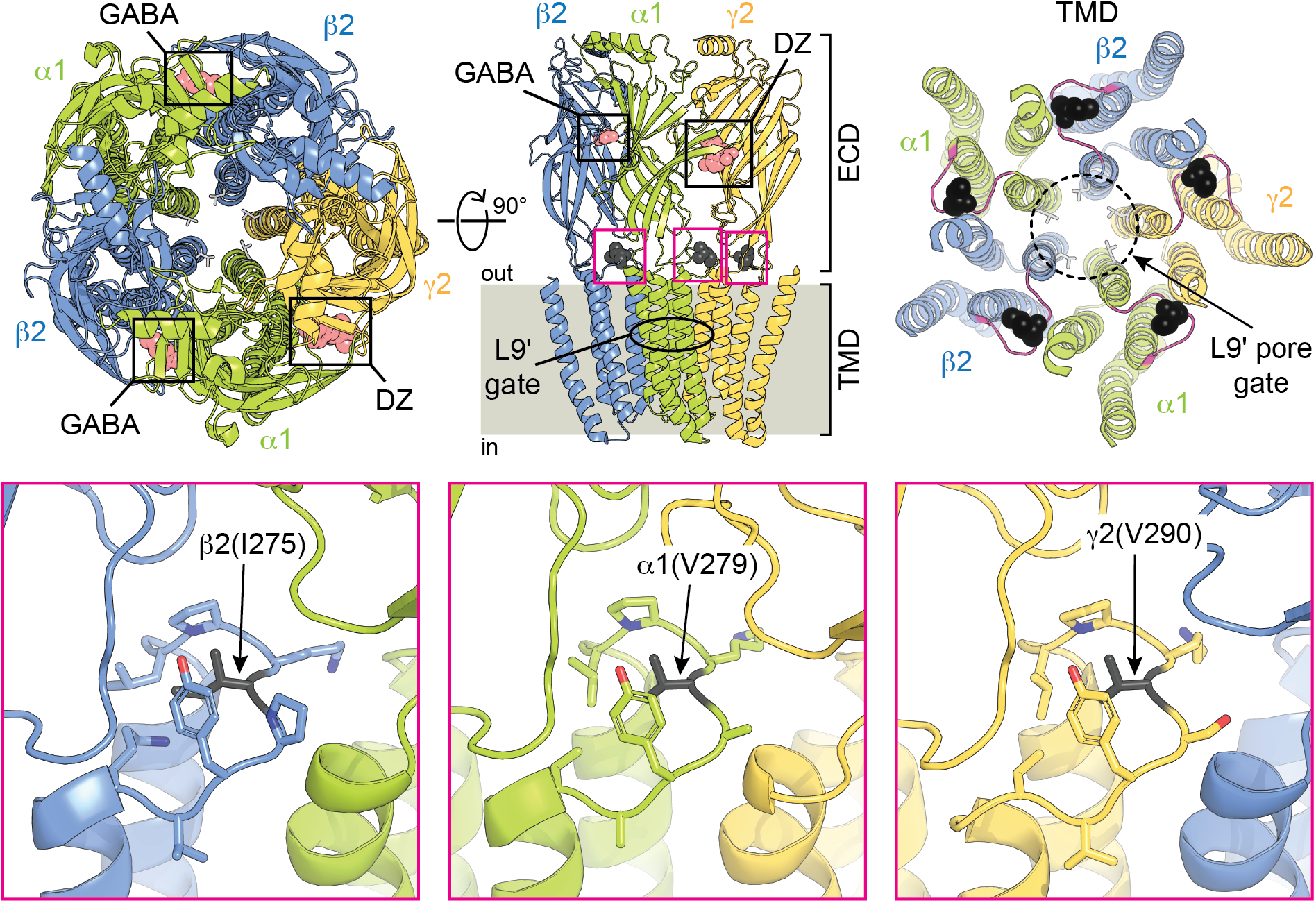
Cryo-EM structural representation of a synaptic GABA_A_α1β2γ2 receptor in complex with GABA and DZ. **(*upper left*)** Top-down view from the extracellular side of the membrane. Subunits α1, β2, and γ2 are shown in green, blue, and yellow, respectively, with bound GABA and DZ as salmon spheres. The 9’ leucines forming the central pore gate are shown as light gray sticks. **(*upper middle*)** Side-on view from the plane of the membrane omitting back two subunits for clarity. The central residue in each M2-M3 linker that was mutated in this study is shown as black spheres. **(*upper right*)** Top-down view with the ECD removed and M2-M3 linkers colored magenta. **(*lower*)** Expanded views of the M2-M3 linker residues that were mutated in this study shown as black sticks. Location of each view corresponds to the magenta boxes in the *upper middle* panel. Cryo-EM map is from PDB 6X3X. Note that rat α1(V279) corresponds to human α1(V280) in the cryo-EM map.

Previously we identified a specific residue, the central valine in the α1 subunit M2-M3 linker (V279), that regulates the efficiency of energetic linkage between DZ bound in the ECD and the pore gate in the TMD (Nors et al., 2021). Both functional and structural observations indicate that channel gating and allosteric modulation by BZDs involves conformational changes throughout the receptor (Baur & Sigel, 2005; Goldschen-Ohm et al., 2010; Sancar & Czajkowski, 2011; Sharkey & Czajkowski, 2008; Venkatachalan & Czajkowski, 2012; Williams & Akabas, 2000) including changes in buried surface area at all inter-subunit interfaces and with the M2-M3 linkers of all subunits undergoing a radial expansion during pore opening (Kim et al., 2020; Masiulis et al., 2019; Zhu et al., 2022). Thus, we investigate here the effects of alanine substitutions in the analogous central position of α1(V279), in the M2-M3 linkers of β2 (I275A) and γ2 (V290A) subunits **(Figure 1)**. Specifically, we explore how these alanine substitutions affect the intrinsic closed-open equilibrium and linkage between DZ bound in the ECD and the pore gate in the TMD. Although DZ does also bind to several lower affinity sites in the TMD (Kim et al., 2020; Masiulis et al., 2019), the observed saturation of DZ responses in this study at 1-3 μM DZ suggests that we are primarily assaying binding to the high affinity site in the ECD (Walters et al., 2000). We show here that alanine substitution at the center of the M2-M3 linker has subunit-specific effects on the intrinsic closed-open equilibrium and its modulation by DZ.

### Rationale for use of a gain-of-function mutant as a background on which to probe the intrinsic pore closed-open equilibrium and linkage between the BZD site and the pore gate

Weak modulatory drugs such as BZDs do not confer appreciable channel opening by themselves, but rather regulate the channel’s response to an agonist such as GABA. Thus, electrophysiological measures of channel current necessitate co-application of both drug and agonist to open the channel. This complicates elucidation of the molecular linkage between the drug site and the pore gate because 1) the agonist’s energetic contribution to pore gating typically dominates as compared to the weak modulator, and 2) it is difficult to distinguish between direct effects of the modulator on the pore equilibrium versus indirect effects via a change in agonist affinity while the modulator is bound. To overcome these challenges and investigate the energetic linkage between the BZD site in the ECD and the pore gate in the TMD, we leverage the gain-of-function mutation α1(L9’T) **(Figure 1)** (Chang & Weiss, 1999; Rüsch & Forman, 2005; Scheller & Forman, 2002). This mutation in the hydrophobic pore gate confers spontaneous exchange between closed and open states in the absence of ligand, to which changes in current in response to either BZDs or mutations can be readily detected. Thus, this mutant allows straightforward quantification of the energetic consequence of BZD binding or a mutation on the pore closed-open equilibrium from conventional measures of channel current (Nors et al., 2021).

Importantly, α1(L9’T)β2γ2 receptors exhibit the same pharmacology as wild type receptors, being blocked by picrotoxin, allosterically modulated by BZDs, and activated robustly by GABA **(Figure 2)** (Rüsch & Forman, 2005; Scheller & Forman, 2002). Kinetic analysis of macroscopic currents elicited with rapid jumps in ligand concentration indicate that nearly all the effects of the α1(L9’T) mutation can be explained by a stabilization of the open conformation with little change to the energetics of closed or desensitized states (Scheller & Forman, 2002). A similar conclusion based on both single channel recordings and macroscopic kinetic analyses was reached for the γ2(L9’S) gain-of-function mutation (Bianchi & Macdonald, 2001). This suggests that the primary effect of the α1(L9’T) mutation is to shift the intrinsic closed-open equilibrium while otherwise maintaining the same mechanism for gating and allosteric modulation as in wild type channels, similar to observations of gain-of-function mutations in nicotinic acetylcholine receptors (Purohit & Auerbach, 2009). If this is true, then activation of wild type channels with lower concentrations of GABA that elicit similar activity to the gain-of-function mutant should be equivalent. However, whenever an agonist such as GABA is used to obtain a desired level of activity, it is difficult to distinguish between a mutation having a direct effect on the pore equilibrium versus an indirect effect via a change in agonist affinity (Colquhoun, 1998). Thus, we chose to use the α1(L9’T) gain-of-function mutation for a uniform background on which to probe the energetics of the pore equilibrium and its linkage with the classical BZD site in the ECD without complication from any changes in agonist affinity. We note that we have both shown previously (Nors et al., 2021) and show here that observations for mutations in the α1(L9’T) background have translated to predicted effects in wild type channels.

**Figure 2.**
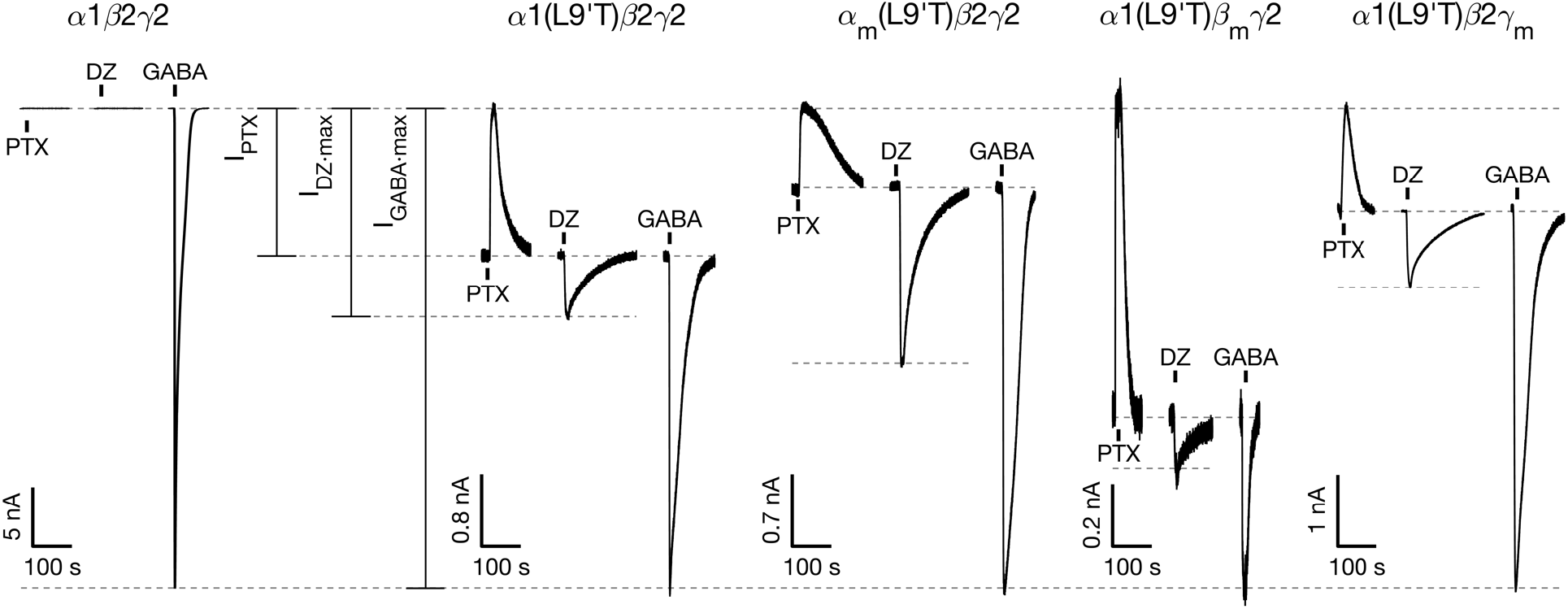
M2-M3 linker mutations α_m_, β_m_, or γ_m_ have subunit-specific effects on the intrinsic closed-open equilibrium and its modulation by DZ. Example currents from wild type α1β2γ2 and gain-of-function α1(L9’T)β2γ2 receptors without and with M2-M3 linker mutations α_m_, β_m_, or γ_m_. Currents are in response to 10 second pulses (black bars) of either 1 mM PTX, saturating 1-3 μM DZ (high affinity ECD site), or saturating GABA. Full concentration-response curves are shown in **Figures S1-S2**. Current block by the pore blocker PTX was used to assess the amount of spontaneous activity and to normalize DZ- and GABA-evoked responses from different oocytes. Currents are normalized from the zero-current baseline in PTX to the maximal GABA-evoked response.

## Results

### Effects of alanine substitutions in the center of the M2-M3 linkers of specific subunits on GABA- and DZ-evoked currents in a gain-of-function receptor

We previously identified a specific residue in the center of the α1 subunit M2-M3 linker (rat V279, human V280) that is involved in mediating linkage between the BZD diazepam (DZ) and the pore gate (Nors et al., 2021). Here, we investigate the effects of alanine substitutions at the analogous central positions in the M2-M3 linkers of rat α1, β2, and γ2 subunits in wild type α1β2γ2 and gain-of-function α1(L9’T)β2γ2 GABA_A_Rs (α_m_ = α1(V279A); β_m_ = β2(I275A); γ_m_ = γ2(V290A)). Note that rat and human sequences are nearly identical, differing by only one residue in α1 and β2 subunits in disordered regions at the N-terminus or in the M3-M4 intracellular linker, respectively, and in γ2 subunits having slightly different lengths of M3-M4 linkers. Xenopus laevis oocytes were co-injected with mRNA for α1, β2, and γ2 subunits (or mutants) in a 1:1:10 ratio (Boileau et al., 2002), and current responses to microfluidic application of ligands were recorded in two-electrode voltage clamp.

For wild type α1β2γ2 receptors, no detectable currents were evoked upon application of the pore blocker picrotoxin (PTX) or the weak modulator DZ, whereas robust currents were elicited with GABA **(Figure 2)**. These observations are as expected for channels that are closed at rest and consistent with DZ being a sufficiently weak modulator that it is unable to open the pore to an observable level on its own. In contrast, receptors harboring the gain-of-function α1(L9’T) mutation exhibited both spontaneous PTX-sensitive current (*I_PTX_*) and additional current beyond the spontaneous current baseline in response to DZ alone (*I_DZ_*) **(Figure 2)**. The ability of DZ to elicit increased channel activity in the gain-of-function mutant parallels its ability to potentiate responses to sub-saturating concentrations of agonist (e.g., GABA) in wild type receptors (Goldschen-Ohm et al., 2014) and illustrates that DZ does energetically bias the pore towards an open conformation. This bias is insufficient to observe channel opening in wild type receptors but is readily detected in the gain-of-function mutant with a much smaller energy difference between closed and open conformations.

For each oocyte we recorded current responses to a series of 10 second pulses of increasing concentrations of either GABA or DZ bookended by 10 second pulses of 1 mM PTX **(Figures S1-2)**. Current block by the pore blocker PTX was used to assess the amount of spontaneous unliganded activity (*I_PTX_*) and to define the zero current baseline. For comparison across oocytes, currents were normalized from the zero current baseline in PTX to the maximal GABA-evoked response (*I_GABA·max_*) **(Figure 2)**. In the absence of a response to saturating GABA to estimate the maximal open probability, DZ-evoked responses (*I_DZ_*) were normalized by matching the PTX-sensitive current amplitude in each DZ recording to the median PTX-sensitive current amplitude across GABA recordings. This amounts to computing the fold-increase in basal open probability conferred by DZ. GABA and DZ concentration-response curves (CRCs) were computed for ligand-evoked current amplitudes with respect to the unliganded current baseline (i.e., the baseline current in the absence of any ligands) and fit to the Hill equation **(Equation 1; Figures S1-8)**. For each construct, the GABA or DZ CRCs were pooled after normalizing to their individual Hill fits, and the pooled data were fit to the Hill equation with *I_max_* = 1.

The gain-of-function mutant α1(L9’T)β2γ2 left-shifts the GABA CRC by ∼100-fold, consistent with previous reports (Rüsch & Forman, 2005) and qualitatively as expected for a bias from closed to open conformations where GABA binds with higher affinity to open versus closed states **(Figure S7)**. In the α1(L9’T)β2γ2 background, the individual mutations α_m_ or γ_m_ had little effect on GABA CRCs, whereas β_m_ right-shifted the CRC by ∼10-fold and conferred a reduced sensitivity with a Hill coefficient less than one **(Figures S4, S7)**. Thus, β_m_ contributes to apparent affinity for GABA binding, whereas α_m_ and γ_m_ do not. None of the individual mutations α_m_, β_m_, or γ_m_ had any effect on DZ CRCs **(Figure S5, S8)**, indicating that the mutations do not appreciably perturb affinity for DZ binding. Reports for the EC_50_ of DZ-potentiation in wild type receptors are comparable to the observed EC_50_ for direct DZ-gating of gain-of-function receptors reported here and in previous studies **(Figure S8)** (Campo-Soria et al., 2006; Li et al., 2013; Nors et al., 2021; Rüsch & Forman, 2005; Walters et al., 2000).

### Asymmetric roles for the central residue of the M2-M3 linker of specific subunits in regulating the pore closed-open equilibrium

The spontaneous unliganded open probability (*P_o·unliganded_*) is given by the product of the maximal open probability in saturating GABA (*P_o·GABA·max_*) and the ratio of the PTX-sensitive to the maximal GABA-Data points are for individual oocytes evoked current amplitudes (*I_PTX_/I_GABA·max_* as shown (a1(L9’T)b2g2: n = 7; a_m_(L9’T)b2g2: n = 4; a1(L9’T)b_m_g2: n = 7; a1(L9’T)b2g_m_: n = 6). Box in **Figure 2**). For wild type α1β2γ2 receptors, *P_o·GABA·max_* ≈ 0.8 (Keramidas & Harrison, 2010). Under the assumption that all gain-of-function constructs (e.g., constructs harboring the α1(L9’T) mutation) should increase *P_o°GABA·max_*, to something approaching one, we estimate for these constructs *P_o°unliganded_* ≈ *I_PTX_*/*I_GABA·max_*. We previously verified from single channel recordings that *P_o°GABA·max_* = 0.93 for αm(L9’T)β2γ2 receptors (Nors et al., 2021). Importantly, our primary conclusions are relatively insensitive to such small errors in the estimation of *P_o°GABA·max_*.

In the α1(L9’T)β2γ2 background, both α_m_ and γ_m_ reduced the unliganded open probability by ∼2-fold **(Figure 3)**. In contrast, β_m_ increased the spontaneous unliganded open probability by ∼2-fold to ∼0.7 which is close to the maximal open probability in saturating GABA for wild type receptors **(Figure 3)**. Thus, mutation of the central residue in the β2 subunit M2-M3 linker from isoleucine to alanine dramatically increases the intrinsic probability of pore opening, whereas the analogous mutations in the α1 or γ2 subunits inhibit spontaneous unliganded activity.

**Figure 3.**
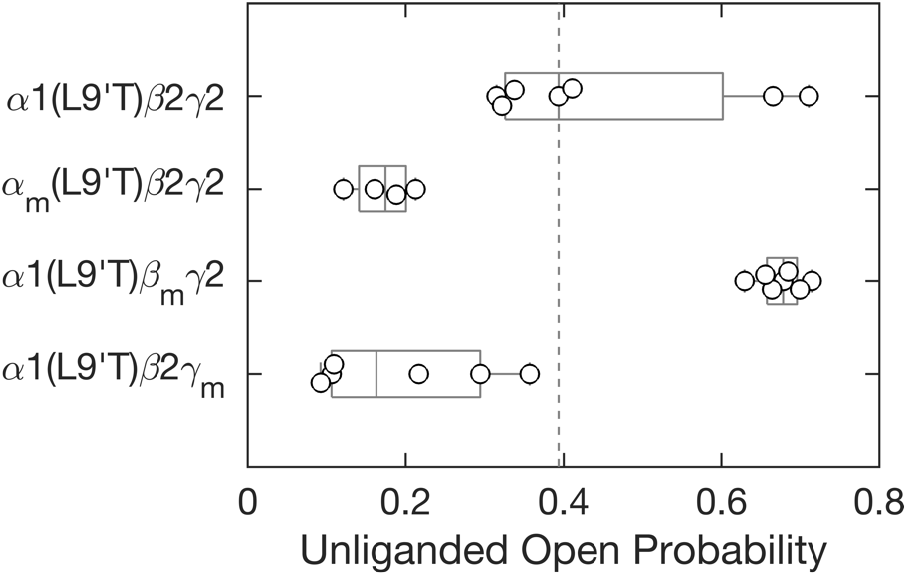
M2-M3 linker mutations either enhance (β_m_) or inhibit (α_m_, γ_m_) intrinsic pore opening in the absence of ligand. Spontaneous open probability for M2-M3 linker mutations α_m_, β_m_, or γ_m_ in the background of the gain-of-function mutation α1(L9’T) estimated as the ratio of PTX-sensitive to maximal GABA-evoked current amplitudes *I_PTX_/I_GABA·max_* (see Figure 2). Data points are for individual oocytes ((α1(L9’T)β2γ2: n = 7; αm(L9’T)β2γ2: n = 4; α1(L9’T)βmγ2: n = 7; α1(L9’T)β2γm: n = 6). Box plots indicate quartiles, and the vertical dashed line is the median for the a1(L9’T)β2γ2 background.

Given the large increase in intrinsic open probability conferred by β_m_ in the α1(L9’T)β2γ2 background, we asked whether β_m_ provides a sufficient bias for an open conformation to detect appreciable unliganded channel opening in the absence of the gain-of-function α1(L9’T) mutation. To test this, we repeated the same PTX and GABA concentration-response protocol described above for α1β_m_γ2 receptors **(Figure 4A, Figure S3)**. The clear PTX-sensitive current verifies that β_m_ alone is sufficient to open the channel pore in the absence of ligand. In contrast, α_m_β2γ2 receptors did not exhibit any obvious PTX-sensitive current, as expected given that α_m_ reduces spontaneous activity in the α1(L9’T)β2γ2 background **(Figure 4A, Figure S3)**. We did not test γ_m_ as it also reduces spontaneous activity similar to α_m_. Consistent with α_m_ inhibiting and β_m_ promoting intrinsic pore opening, in the wild type α1β2γ2 background α_m_ or β_m_ right- or left-shifted GABA CRCs, respectively **(Figures S6-7)**. Given that β_m_ is a gain-of-function mutation, we estimated the unliganded open probability in the same manner as for channels with the α1(L9’T) mutation. In comparison to α1(L9’T)β2γ2, α1β_m_γ2 receptors were open spontaneously approximately half as much, whereas the combination of both mutations in α1(L9’T)β_m_γ2 receptors conferred the most unliganded channel opening, consistent with effects of the gain-of-function mutations α1(L9’T) and β_m_ being at least partially additive and independent **(Figure 4B)**. However, see below in the section on combinations of mutations for their non-additive energetic effects on the intrinsic closed-open equilibrium.

**Figure 4.**
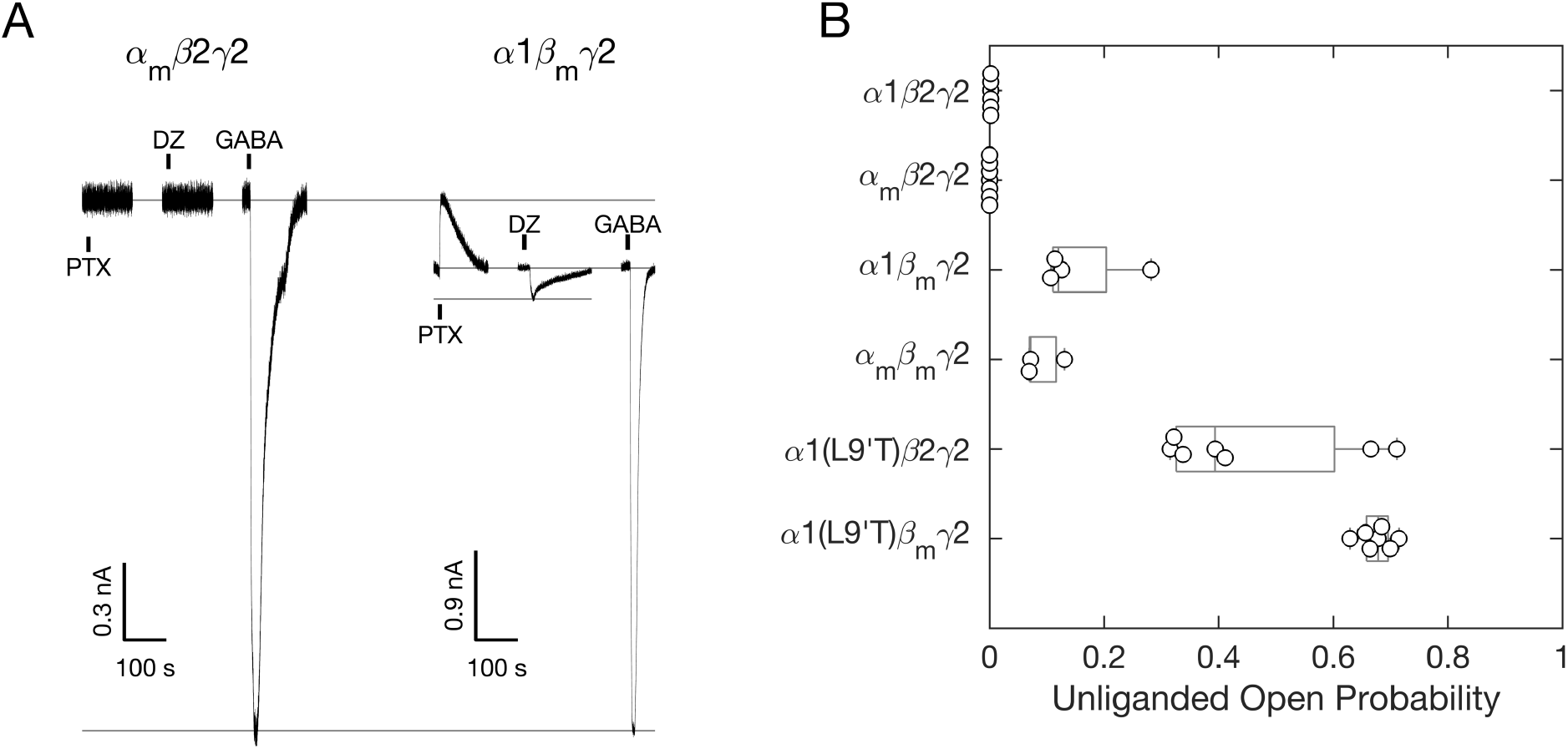
The central residue (I275) in the β2 subunit M2-M3 linker is required to keep the channel closed in the absence of ligand. **(A)** Example currents from receptors with α_m_ or β_m_ mutations in a wild type α1β2γ2 background. Currents are in response to 10 second pulses (black bars) of either 1 mM PTX, saturating 3 μM DZ (high affinity ECD site), or saturating GABA. Full concentration-response curves are shown in **Figure S3**. Current block by the pore blocker PTX was used to assess the amount of spontaneous activity and to normalize DZ- and GABA-evoked responses from different oocytes. Currents are normalized from the zero-current baseline in PTX to the maximal GABA-evoked response. **(B)** Spontaneous unliganded open probability for M2-M3 linker mutations α_m_ and/or β_m_ in wild type α1β2γ2 and gain-of-function α1(L9’T)β2γ2 backgrounds estimated as the ratio of PTX-sensitive to maximal GABA-evoked current amplitudes *I_PTX_/I_GABA·max_* (see **(A)** and Figure 2). Data points for constructs including the α1(L9’T) mutation are as shown in Figure 3. Data points are for individual oocytes (α1β2γ2: n = 6; α_m_β2γ2: n = 7; α1β_m_γ2: n = 4; α_m_β_m_γ2: n = 3; α1(L9’T)β2γ2: n = 7; α1(L9’T)β_m_γ2: n = 7). Box plots indicate quartiles.

These data show that the central position in the β2 subunit M2-M3 linker is important for regulating the intrinsic closed-open equilibrium in GABA_A_ receptors. Unlike α1(L9’T), β_m_ (I275A) is located outside of the pore, suggesting that this regulation may overlap with events underlying the transduction of the chemical energy from ligand binding to pore gating. In contrast, alanine substitutions at the analogous position in the M2-M3 linker of α1 or γ2 inhibit pore opening, indicating an asymmetric role for specific subunit M2-M3 linkers in regulating the pore closed-open equilibrium.

### Asymmetric roles for the central residue of the M2-M3 linker of specific subunits in regulating the efficiency of DZ-to-pore linkage

Despite having little effect on DZ apparent affinity, we previously identified that the α_m_ mutation increases the efficiency of transduction of chemical energy from DZ binding to gating of the channel pore by up to 3-fold (Nors et al., 2021). Intriguingly, this increase in DZ efficiency was only observed for alanine substitution at the central residue (V279) of the rat α1 subunit M2-M3 linker, whereas alanine substitutions at other residues in the linker had no effect (Nors et al., 2021). In addition to its high affinity site in the ECD **(Figure 1)**, DZ also binds to several lower affinity sites in the TMD (Kim et al., 2020; Masiulis et al., 2019). However, the observed saturation of DZ responses in this study at 1-3 μM DZ suggests that we are primarily assaying binding to the high affinity site in the ECD (Walters et al., 2000). Thus, we interpret our results as probing linkage between the high affinity BZD site in the ECD and the pore gate in the TMD.

Given that structural changes conferred upon DZ binding appear to involve global changes in conformation with similar gross overall changes in each subunit (Kim et al., 2020; Masiulis et al., 2019), as well as functional evidence that BZDs confer global conformational changes at multiple subunit-subunit interfaces (Baur & Sigel, 2005; Goldschen-Ohm et al., 2010; Sancar & Czajkowski, 2011; Sharkey & Czajkowski, 2008; Venkatachalan & Czajkowski, 2012; Williams & Akabas, 2000), we asked whether the analogous alanine substitutions at the center of the M2-M3 linkers in β2 or γ2 subunits would confer similar effects on DZ efficiency. We employed a simple channel gating scheme between closed (C) and open (O) pore states in both unliganded and DZ-bound conditions to quantify the energetic linkage between DZ binding and pore gating **(Figure 5A)**. In the absence or presence of DZ we estimated the maximal unliganded or DZ-evoked open probability as either *P_o·unliganded_* = *I_PTX_/I_GABA·max_* or *P_o·DZ·bound_* = *I_DZ·max_/I_GABA·max_* (see **Figure 2**). The free energy difference between closed and open states in unliganded (Δ*G_unliganded_*) or DZ-bound (Δ*G_DZ·bound_*) conditions is given by **Equations 2-3**. The energetic consequence of DZ binding on the pore closed-open equilibrium is the difference between DZ-bound and unliganded conditions: ΔΔ*G_DZ_* = Δ*G_DZ·bound_* − Δ*G_unliganded_*.

**Figure 5.**
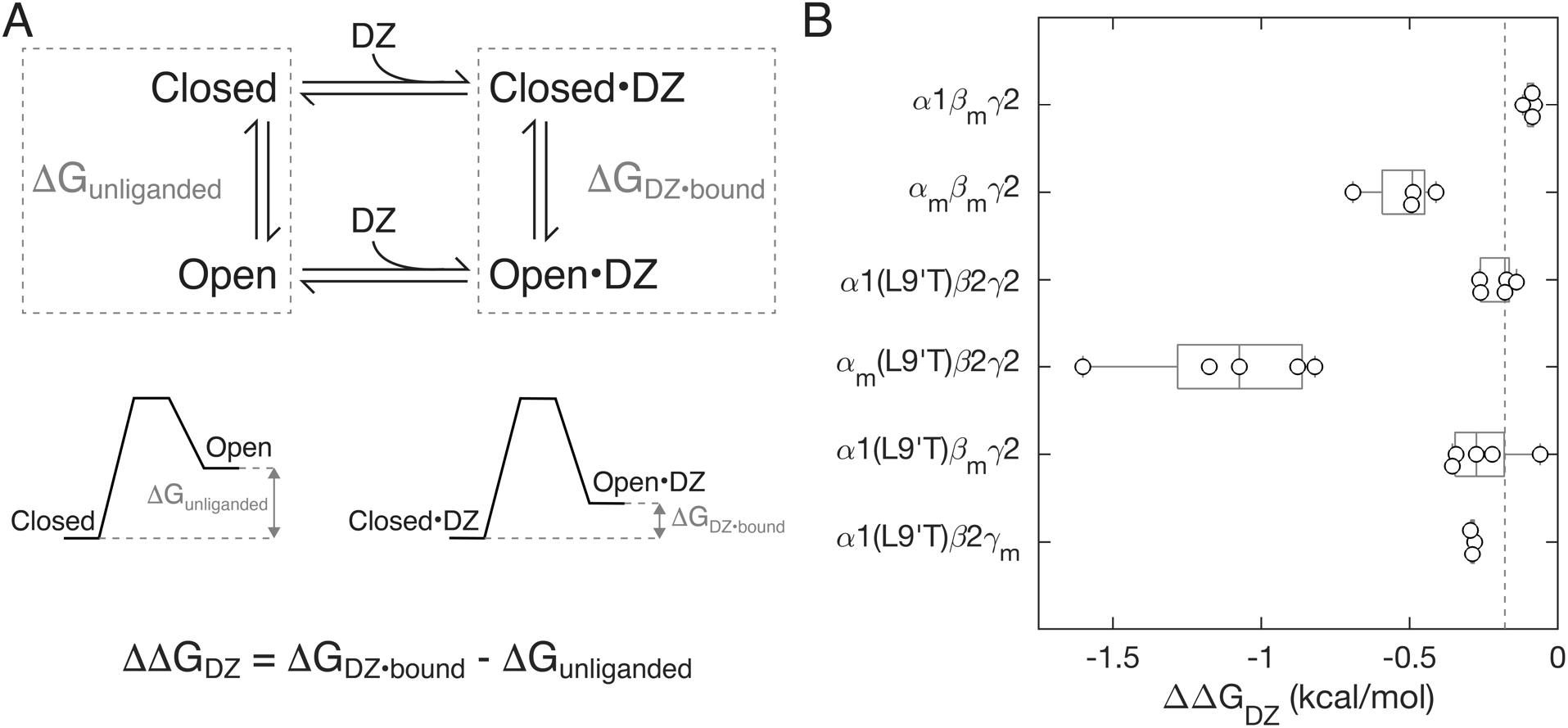
The α1 subunit M2-M3 linker mutation α_m_ enhances DZ modulation of pore gating, whereas analogous mutations in β2 or γ2 subunits have little effect. **(A)** Simple scheme for gating between closed and open conformations (vertical arrows) with DZ binding and unbinding (horizontal arrows) separating unliganded and DZ-bound states. The free energy difference between open and closed conformations in both unliganded (Δ*G_unliganded_*) and DZ-bound (Δ*D_DZ·bound_*) conditions are depicted graphically below the scheme. See **Equations 2-3** for how these energies are computed. **(B)** The change in closed versus open free energies upon DZ binding in the ECD (see **Equation 4**) for α_m_, β_m_, or γ_m_ mutations in wild type α1β2γ2 and/or gain-of-function α1(L9’T)β2γ2 backgrounds. The more negative the value of ΔΔG_DZ_, the more DZ increases channel open probability. Data points are for individual oocytes (α1β_m_γ2: n = 4; α_m_β_m_γ2: n = 4; α1(L9’T)β2γ2: n = 5; α_m_(L9’T)β2γ2: n = 5; α1(L9’T)β_m_γ2: n = 5; α1(L9’T)β2γ_m_: n = 3). Box plots indicate quartiles, and the vertical dashed line is the median for the α1(L9’T)β2γ2 background.

In contrast to our prior observation that α_m_ increases the energetic linkage between DZ binding and pore gating (Nors et al., 2021), we observe little to no effect of the analogous mutations β_m_ or γ_m_ on ΔΔ*G_DZ_* **(Figure 5B)**. These data suggest that linkage between DZ-binding and the pore gate is mediated differentially by M2-M3 linkers of specific subunits.

### Combinations of M2-M3 linker mutations with subunit-specific effects

To assess the independence of α_m_, β_m_, and γ_m_ mutations we measured GABA and DZ CRCs, spontaneous unliganded open probabilities, and DZ-to-pore linkage energetics for combinations of mutations in the α1β2γ2 and α1(L9’T)β2γ2 backgrounds in the same manner as for the individual mutations. In the α1(L9’T)β2γ2 background, all double mutants and the triple mutant right-shifted GABA CRCs by ∼3- to 10-fold with either similar or increased sensitivities, i.e., Hill coefficients **(Figures S4, S7)**. Interestingly, every subunit combination that includes the β_m_ mutation exhibits a similar GABA EC_50_ independent of whether the α1(L9’T) mutation is present. If the shifts in GABA EC_50_ reflect only the degree of bias toward the higher affinity open conformation, then the combination of α1(L9’T) and β_m_ should be even more left-shifted than either mutation alone. However, we observe a right-shift in GABA EC_50_ for α1(L9’T)β_m_γ2 as compared to α1(L9’T)β2γ2, suggesting that β_m_ has a dominant effect on setting the affinity for GABA binding in the extracellular domain in addition to its effect on the intrinsic pore equilibrium. In contrast, nearly all combinations of α_m_, β_m_, and γ_m_ mutations had little effect on DZ CRCs, like observations for the individual mutations, the exception being a 4- fold right-shift in the DZ EC_50_ for α_m_β_m_γ2 **(Figures S5, S6B, S8)**.

For nearly all tested combinations of mutations the intrinsic probability of spontaneous unliganded pore opening was enhanced by β_m_ and inhibited by α_m_ and/or γ_m_ **(Figure 6)**. The only exception is that α1(L9’T)β_m_γ_m_ receptors exhibit similar or even more spontaneous activity than α1(L9’T)β_m_γ2 receptors, despite the addition of γ_m_. The reason for this exception is not clear. The combination of β_m_ and α1(L9’T) conferred a spontaneous unliganded open probability that was roughly the sum of the spontaneous open probabilities for each of the individual mutations. However, the effect of the individual mutations β_m_ and α1(L9’T) on the free energy difference between closed and open states in the absence of ligand (Δ*G_unliganded_*) are not additive. For wild type α1β2γ2 receptors we estimate *P_o·unliganded_* = 0.002 (Mortensen et al., 2003), from which **Equation 2** implies Δ*G_unliganded_* = 3.7 *kcal/mol*. Either β_m_ or α1(L9’T) reduce Δ*G_unliganded_* to 1.0 or 0.1 kcal/mol, respectively, thereby increasing the probability of spontaneous channel opening. However, for the combination of both mutations in α1(L9’T)β_m_γ2 receptors Δ*G_unliganded_* = −0.3 *kcal/mol*, which is only slightly less than for α1(L9’T) alone. Thus, the effects of β_m_ outside of the pore and α1(L9’T) at the pore gate on the channel’s closed-open equilibrium are energetically non-independent. It seems plausible that both mutations may promote similar conformational changes to enhance channel opening.

**Figure 6.**
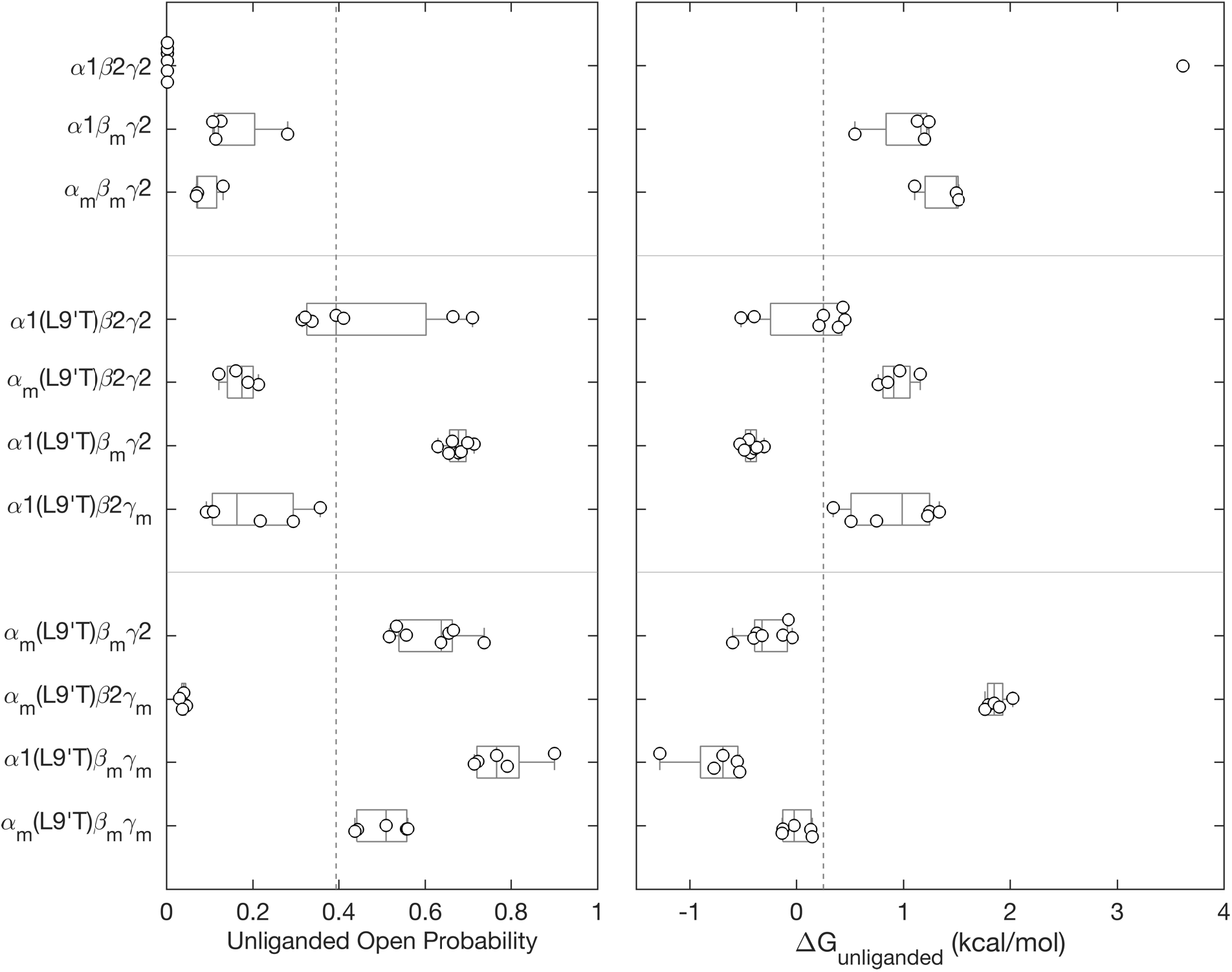
M2-M3 linker mutations either enhance (β_m_) or inhibit (α_m_, γ_m_) intrinsic pore opening in the absence of ligand. **(*left*)** Spontaneous unliganded open probability for receptors with M2-M3 linker mutations α_m_, β_m_, and/or γ _m_ in the gain-of-function α1(L9’T)β2γ2 or wild type α1β2γ2 backgrounds estimated as the ratio of PTX-sensitive to maximal GABA-evoked current amplitudes (see **Figures 2, 4**). **(*right*)** Free energy difference from closed to open states calculated from the unliganded open probability using **Equation 2**. Δ*_o·unliganded_* for α1β2γ2 receptors is based on the estimate *P_o.unliganed_* = 0.002 (Mortensen et al., 2003) which is undetectable in our assay. All other data points are for individual oocytes (α1β2γ2: n = 6; α1β_m_γ2: n = 4; α_m_β_m_γ2: n = 3; α1(L9’T)β2γ2: n = 7; α_m_(L9’T)β2γ2: n = 4; α1(L9’T)β_m_γ2: n = 7; α1(L9’T)β2γ_m_: n = 6; α_m_(L9’T)β_m_γ2: n = 7; α_m_(L9’T)β2γ_m_: n = 5; α1(L9’T)β_m_γ_m_: n = 5; α_m_(L9’T)β_m_γ_m_: n = 5). Box plots indicate quartiles, and the vertical dashed line is the median for the α1(L9’T)β2γ2 background. The unliganded open probabilities for some of the constructs are the same as shown in Figures 3-4.

In the α1(L9’T)β2γ2 background the combination of α_m_ and γ_m_ on Δ*G_unliganded_* was roughly the sum of the effects of the individual mutations **(Figure 6)**. Thus, inhibition of pore opening by α_m_ and γ_m_ is largely independent. In contrast, Δ*G_unliganded_* for combinations of either α_m_ or γ_m_ with β_m_ was similar to that for β_m_ alone. Thus, the enhancement of channel gating conferred by β_m_ largely dominates such that α_m_ or γ_m_ can no longer confer their inhibitory effects. Thus, whereas α_m_ and γ_m_ are largely independent of each other, they are not independent of β_m_. However, in the triple mutant α_m_(L9’T)β_m_γ_m_ the combination of both α_m_ and γ_m_ is sufficient to slightly inhibit the gain-of-function conferred by β_m_.

The combined effect of combinations of α_m_ with β_m_ and/or γ_m_ mutations on DZ-to-pore linkage ΔΔ*G_DZ_* were uniformly less than additive, suggesting that the mutations are not completely independent **(Figure 7)**. Nonetheless, the qualitative observation that α_m_ enhances the efficiency of DZ-gating remains true when in combination with either β_m_ or γ_m_ in both α1β2γ2 and α1(L9’T)β2γ2 backgrounds. The enhancement is reduced when α_m_ is combined with either β_m_ or γ_m_, and reduced further for the combination of all three mutations α_m_, β_m_, and γ_m_. The mechanism for the non-additive effects of combinations of mutations on ΔΔ*G_DZ_* is unclear but indicates that it is at least possible for M2-M3 linkers from each subunit to contribute to DZ-to-pore linkage.

**Figure 7.**
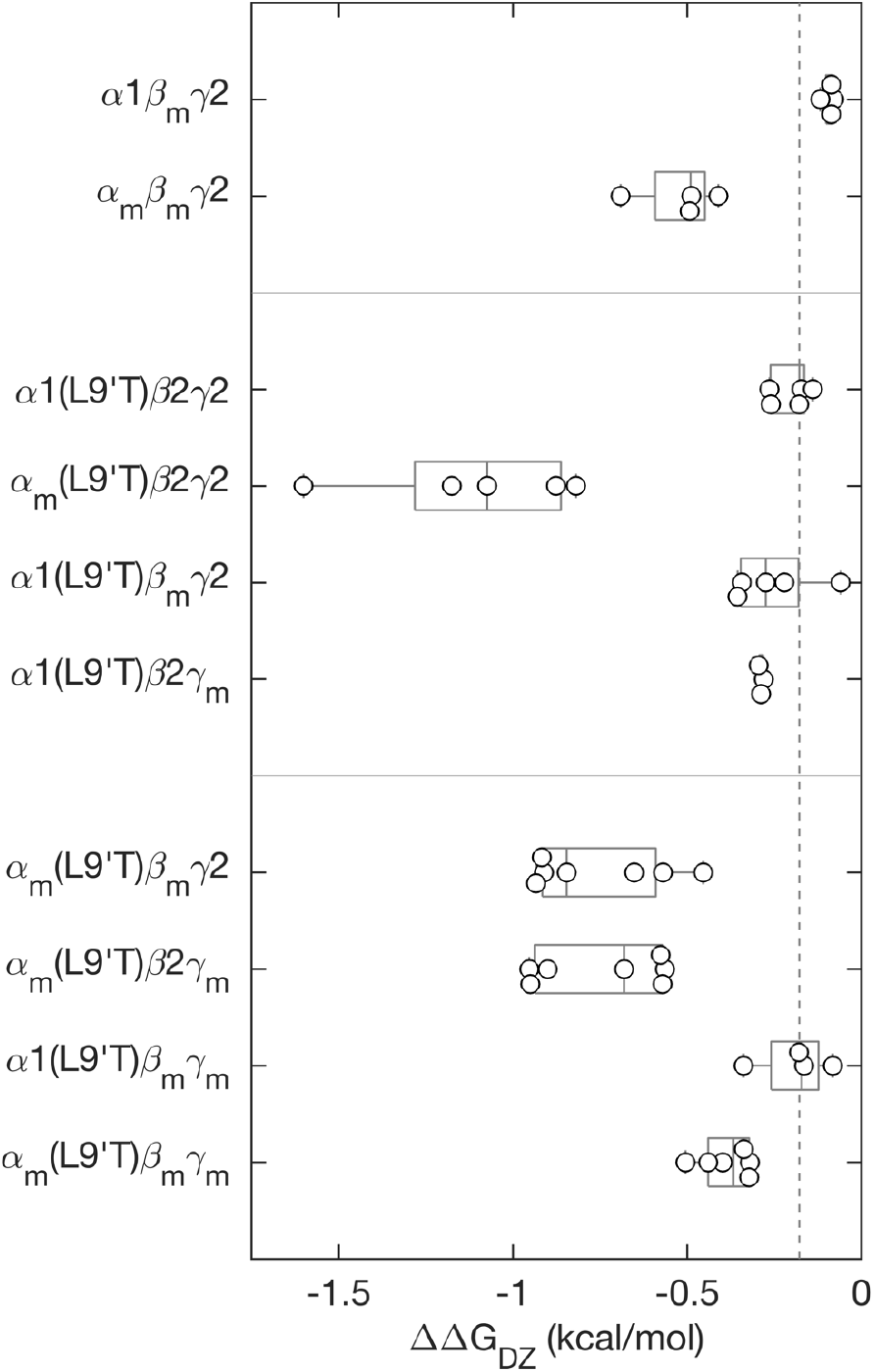
M2-M3 linker mutations either enhance (α_m_) or have no effect (β_m_, γ_m_) on the energy DZ-binding in the ECD provides to the channel’s closed-open equilibrium. The change in the free energy difference from closed to open states upon DZ binding in the ECD (see **Equation 4**) for α_m_, β_m_, or γ_m_ mutations in wild type α1β2γ2 and/or gain-of-function α1(L9’T)β2γ2 backgrounds. The more negative the value of ΔΔG_DZ_, the more DZ increases channel open probability. Data points are for individual oocytes (α1β_m_γ2: n = 4; α_m_β_m_γ2: n = 4; α1(L9’T)β2γ2: n = 5; α_m_(L9’T)β2γ2: n = 5; α1(L9’T)β_m_γ2: n = 5; α1(L9’T)β2γ_m_: n = 3; α_m_(L9’T)β_m_γ2: n = 7; α_m_(L9’T)β2γ_m_: n = 7; α1(L9’T)β_m_γ_m_: n = 4; α_m_(L9’T)β_m_γ_m_: n = 6). Box plots indicate quartiles, and the vertical dashed line is the median for the α1(L9’T)β2γ2 background. The data for some of the constructs are the same as shown in Figure 5B.

## Discussion

We show that alanine substitutions at the central residue in the M2-M3 linkers of specific subunits α_m_, β_m_ or γ_m_ have asymmetric effects on the intrinsic unliganded closed-open equilibrium and its modulation by DZ. Whereas α_m_ and γ_m_ inhibit pore opening, β_m_ promotes opening to the extent that this mutation alone is sufficient to cause the channel to open spontaneously in the absence of agonist. Estimating the unliganded open probability in wild type α1β2γ2 receptors as 0.002 (Mortensen et al., 2003), the energetic effect of β_m_ on the closed-open equilibrium is -2.5 kcal/mol (using **Equation 2**), which is more than half the -4.5 kcal/mol supplied by GABA binding to the two neurotransmitter sites (i.e., for a change in open probability from 0.002 to 0.8). In contrast, only α_m_ increases the efficiency of DZ-to-pore linkage, whereas β_m_ or γ_m_ have no effect.

Comparison of GABA_A_R and other pLGIC structures in closed antagonist-bound and activated/desensitized agonist-bound conformations suggests that gating involves grossly symmetric motions of all five subunits including a radial expansion of the M2 pore lining helices and M2-M3 linkers (Gibbs et al., 2023; Nemecz et al., 2016). Similarly, comparison of GABA_A_R structures with and without BZDs suggest that BZDs such as DZ confer a global compaction of the ECD with increased intersubunit contacts, although the observed conformational changes are not large and differ in magnitude based on the solubilization strategy (Kim et al., 2020; Masiulis et al., 2019). However, all structures with bound BZD to-date were obtained in the presence of bound GABA where the complex with GABA is expected to energetically dominate the observed conformation. As discussed above, GABA binding to both agonist sites confer approximately -4.5 kcal/mol to channel gating as compared to -0.4 kcal/mol for DZ in the ECD site (Nors et al., 2021). Furthermore, the observed effect of DZ decreases with increasing baseline activity, similar to recent observations for the energetic effect of the anesthestic propofol and neurosteroid etiocholanolone on the closed/open equilibrium (Pierce et al., 2023). Indeed, functional studies show that DZ fails to potentiate responses to saturating GABA (Lavoie & Twyman, 1996; Perrais & Ropert, 1999; Rogers et al., 1994; Twyman et al., 1989; Vicini et al., 1987), suggesting that the presence of GABA may occlude the conformational effects of DZ in these structures. Nonetheless, there is also functional evidence for more global effects of BZDs on intersubunit interfaces throughout the receptor (Baur & Sigel, 2005; Goldschen-Ohm et al., 2010; Sancar & Czajkowski, 2011; Sharkey & Czajkowski, 2008; Venkatachalan & Czajkowski, 2012; Williams & Akabas, 2000). Indeed, the non-additive effects of various combinations of the mutations examined here indicate some cooperativity between the M2-M3 linkers of all subunits consistent with a global conformational change. However, the effects of individual mutations α_m_, β_m_ or γ_m_ are highly asymmetric.

We hypothesize that the decreased sidechain volume upon substitution of the central linker residue with alanine (α_m_, β_m_, γ_m_) removes steric packing in the center of the roughly arc-shaped structure that the linker adopts and thereby confers an increase in linker flexibility **(Figure 1)**. Our prior observation that neither α1(V297W) nor α1(V297D) enhance DZ-to-pore linkage as does α1(V297A) is consistent with this idea (Nors et al., 2021). However, it is unclear as to why increased flexibility in the M2-M3 linker of different subunits confers such disparate effects. A cryo-EM structural model of α1β3γ2 receptors suggests that the domain including the M2-M3 linker and top of the M2 and M3 helices may be intrinsically more flexible in β2/3 subunits than in α1 or γ2 subunits due to differential stabilization via intrasubunit interactions with a conserved 19’ arginine near the top of the M2 helix (Masiulis et al., 2019). A similar orientation of R19’ in one of the β subunits as compared to the other subunits was found in a recent structure of a native receptor from mouse brain (Sun et al., 2023). Changes in linker flexibility and intersubunit contacts have also been associated with closed versus open conformations of glycine receptors (Du et al., 2015). Thus, a difference in intrinsic flexibility may contribute to the disparate effects of α_m_, β_m_ or γ_m_ on the closed-open equilibrium. However, the opposing effects of β_m_ as compared to α_m_ or γ_m_ on pore opening suggests that linker flexibility does not automatically map to increased channel activity but depends on the intersubunit interface in which the linker is located.

Changes in M2-M3 linker flexibility could also alter the overall compaction of the TMD helices thought to affect coupling with the BZD site (Kim et al., 2020). The observation that only α_m_ enhances the efficiency of DZ-to-pore linkage whereas β_m_ or γ_m_ does not suggest that the effects of the mutations on TMD compaction are either 1) global but opposing, 2) more localized to their respective intersubunit interfaces, or 3) not relevant. The idea of distinct effects at specific subunit-subunit interfaces is interesting given that the M2-M3 linkers in β subunits are located at the β/α intersubunit interfaces below the agonist binding sites, whereas the M2-M3 linker in one of the α subunits is located at the α/γ intersubunit interface below the BZD binding site **(Figure 8)**. Thus, physical location of each M2-M3 linker with respect to either agonist or BZD binding sites in the ECD may explain the subunit-specific effects of α_m_, β_m_ or γ_m_ on the intrinsic closed-open equilibrium and its modulation by DZ. In such a case the M2-M3 linkers of distinct subunits would have asymmetric roles in pore gating and drug modulation, with the β subunit M2-M3 linker having a predominant role in regulating pore opening and the α subunit M2-M3 being most involved in DZ-modulation.

**Figure 8.**
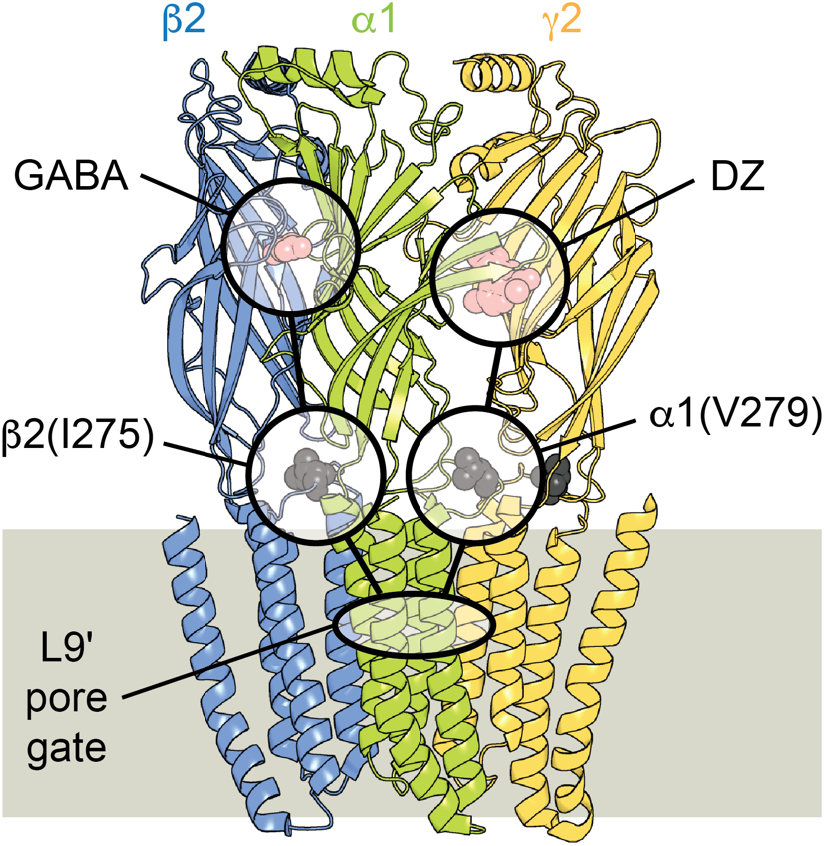
Functional asymmetry in M2-M3 linkers may correspond to the ligand binding interface in which the linker is located. Side-on view from the plane of the membrane omitting back two subunits for clarity. Subunits a1, b2, and g2 are shown in green, blue, and yellow, respectively. The relative locations of bound GABA or DZ and M2-M3 linker residues b2(I275) or a1(V279) at β/α or α/γ intersubunit interfaces, respectively, and the 9’ leucine pore gate are highlighted.

Although grossly symmetric, comparison of structures of α1β2/3γ2 receptors in closed antagonist-bound and desensitized agonist-bound conformations indicates some asymmetry in subunit motions. For example, the ECDs of β subunits rotate a few degrees further than other subunits, the M2-M3 linkers of β subunits undergo the largest radial expansion, and the pore gate 9’ leucine in β subunits rotate furthest out of the ion conducting pathway **(Figure S9)** (Kim et al., 2020; Masiulis et al., 2019). Observations for disulfide bond formation and zinc binding to introduced cysteines also suggest asymmetric flexibility between α and β subunits near the top of the M2 helix (Horenstein et al., 2005). From a functional perspective, mutation of a conserved lysine in the M2-M3 linker that is associated with human epilepsy (α1(K278M), β2(K274M), γ2(K289M)) has subunit-specific effects on receptor activity (Baulac et al., 2001; Hales et al., 2006). Thus, the asymmetric effects of alanine substitutions in distinct M2-M3 linkers on pore gating and DZ modulation that we observe here may reflect distinct conformational changes in specific subunits or at specific intersubunit interfaces. However, our observations only suggest that the subunits contribute to the energetics of these processes differentially. Global conformational changes that are grossly symmetric amongst all subunits, but for which distinct subunit M2-M3 linkers contribute differentially to the energetics, is entirely compatible with our results.

Here, we show that the central residue in the M2-M3 linkers of β2 versus α1 and γ2 subunits have opposing roles in regulating the intrinsic unliganded closed-open equilibrium. In contrast, only this position in the α1 subunit regulates the efficiency of modulation of this equilibrium by DZ. These observations shed new light on the subunit-specific roles of M2-M3 linkers which correlate structurally with their respective ligand-binding intersubunit interfaces.

## Methods

### Mutagenesis and expression in oocytes

DNA for rat GABA_A_R α1, β2, and γ2 subunits were a gift from Dr. Cynthia Czajkowski. Note that the long isoform of γ2 was used throughout. The α1(L9’T) and α_m_ mutations were introduced individually or serially by site-specific mutagenesis (QuikChange II, Qiagen). Mutations β_m_ and γ_m_ were obtained from GENEWIZ^TM^. Each construct was verified by forward and reverse sequencing of the entire gene. For expression in Xenopus laevis oocytes (EcoCyte Bioscience, Austin, TX) cRNA for each construct was generated from DNA plasmids (mMessage mMachine T7, Ambion). Oocytes were injected with 27–54 ng of total mRNA for α, β, and γ subunits (or mutants) in a 1:1:10 ratio (Boileau et al., 2002) (Nanoject, Drummond Scientific). Oocytes were incubated in ND96 (in mM: 96 NaCl, 2 KCl, 1 MgCl2, 1.8 CaCl2, 5 HEPES, pH 7.2) with 100 mg/ml gentamicin at 18°C.

### Two electrode voltage clamp recording and analysis

Currents from expressed channels 1–3 days post-injection were recorded in two-electrode voltage clamp (Dagan TEV-200 amplifier, HEKA ITC digitizer and Patchmaster software). Oocytes were held at -80 mV and perfused continuously with buffer (ND96) or buffer containing PTX, GABA, or DZ. PTX was diluted from a 1 M stock solution in DMSO. DZ was diluted from a 10 mM stock solution in DMSO. Fresh PTX and DZ stock solutions were tested several times with no change in results. GABA was dissolved directly from powder. A microfluidic pump (Elveflow OB1 MK3+) and rotary valve (Elveflow MUX Distributor) provided consistent and repeatable perfusion and solution exchange across experiments, which limited solution exchange variability to primarily differences between oocytes only. Ten second pulses of PTX, GABA, or DZ were followed by 3–6 min in buffer to allow currents to return to baseline. Recorded currents were analyzed with custom scripts in MATLAB (Mathworks).

Recordings of concentration–response relationships were bookended by pulses of PTX to correct for any drift or rundown during the experiment and to identify the zero current baseline. As described previously (Nors et al., 2021), in some oocytes we accounted for a gradual rundown of channel current during the time course of the recording by applying a linear scaling in time so that the initial and final PTX responses were of equal amplitude. It is worth noting that such scaling had little to no effect on the ratio of the maximal GABA- or DZ-evoked responses (*I_PTX_/I_GABA·max_* or *I_DZ·max_*) to the following final PTX response (*I_PTX_*) as used to estimate open probability and conferred only minor shifts at most to the concentration-response relations. The amount of current rundown was variable across oocytes, with no clear relation to specific constructs.

Current (*I*) concentration–response curves (CRCs) were measured with respect to the unliganded current baseline and fit to the Hill equation:

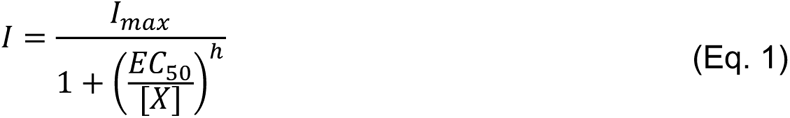

where *I_max_* is the peak current response relative to the unliganded current baseline, [*X*] is ligand concentration, EC_50_ is the concentration eliciting a half-maximal response, and *h* is the Hill coefficient. Note that although the parameters from Hill fits provide a general metric for assessing apparent affinity and sensitivity, they are not easily translatable into physical parameters such as affinities of specific sites or numbers of bound ligands (Holt & Ackers, 2009; Prinz, 2010).

### DZ-gating model

The DZ-gating model **(Figure 5A)** is the same as described previously (Nors et al., 2021). Briefly, the free energy difference from closed to open states was calculated for unliganded (Δ*G_unliganded_*) or DZ-bound (Δ*G_DZ·bound_*) receptors as follows:

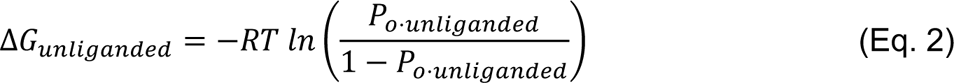

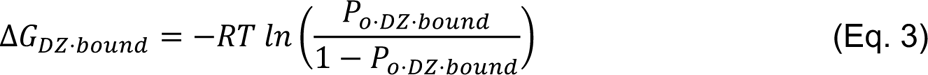

where *RT* is the product of the gas constant and temperature, and we estimated *P_o·unliganded_* ≈ *I_PTX_*/*I_PTX_/I_GABA·max_* and *P_o·DZ·bound_* ≈ *I_DZ·max_*/*I_PTX_/I_GABA·max_*. This estimate is reasonable for gain-of-function mutants whose open probability in saturating GABA is close to one. See **Figure 2** for an illustration of *I_PTX_*, *I_DZ·max_*, and *I_PTX_/I_GABA·max_*. The energetic consequence of DZ binding on the pore closed-open equilibrium ΔΔ*G_DZ_* is the difference between DZ-bound and unliganded conditions:

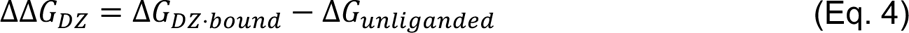

