## Supplemental Figures for "Asymmetric roles for M2-M3 linkers of specific GABA_A_ receptor subunits in the intrinsic closed-open equilibrium and its modulation by the benzodiazepine diazepam"

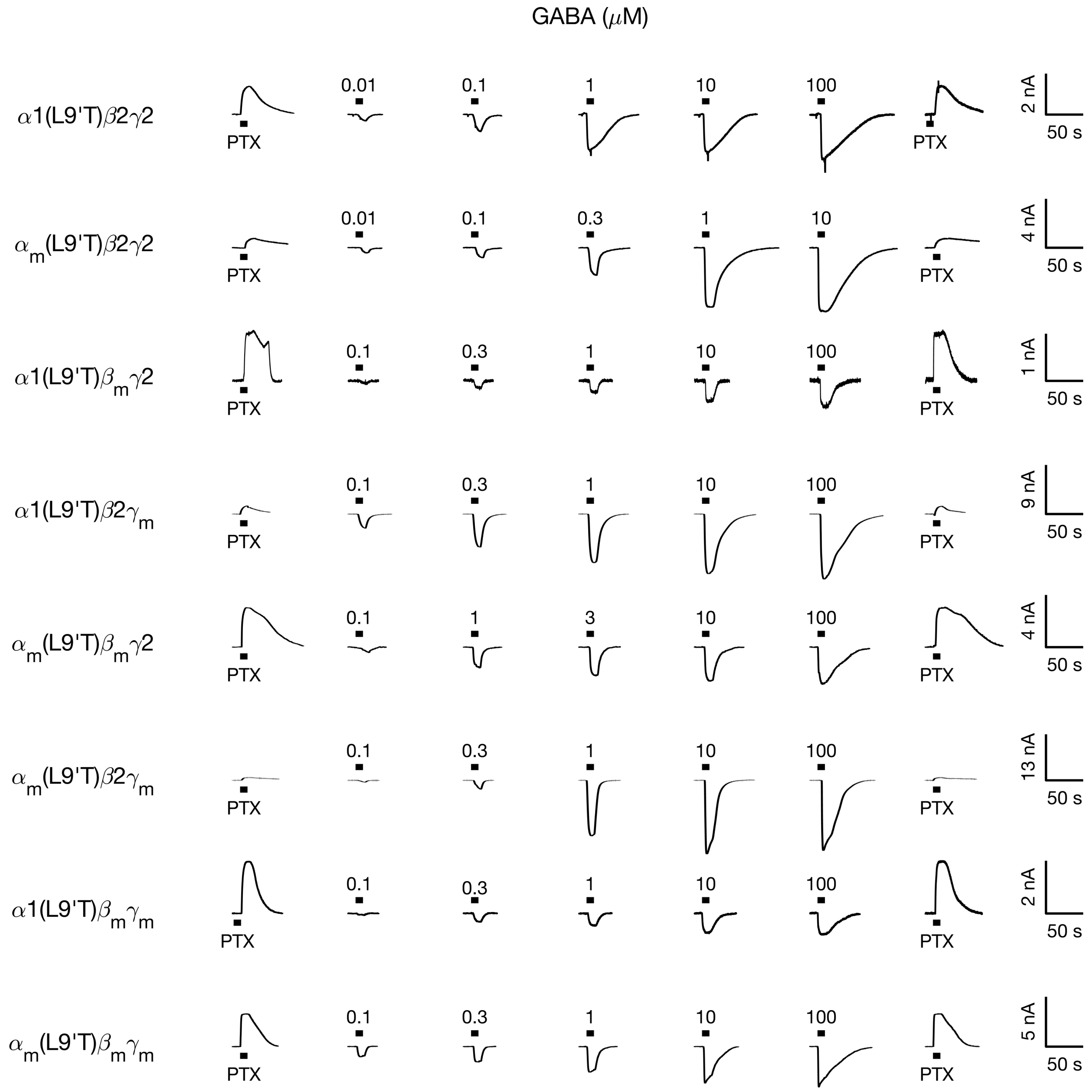


**Figure S1. Example GABA-evoked currents from receptors with M2-M3 linker mutations $\alpha_m$, $\beta_m$, and/or $\gamma_m$ in the gain-of-function $\alpha$1(L9'T)$\beta$2$\gamma$2 background.** Currents are in response to 10 second pulses (black bars) of either 1 mM PTX or a series of increasing concentrations of GABA (concentration in micromolar above respective black bar). Summaries of concentration-response curves across oocytes are shown in **Figure S4**. Responses to GABA were bookended by the pore blocker PTX to assess the amount of spontaneous unliganded activity and identify the zero-current baseline.

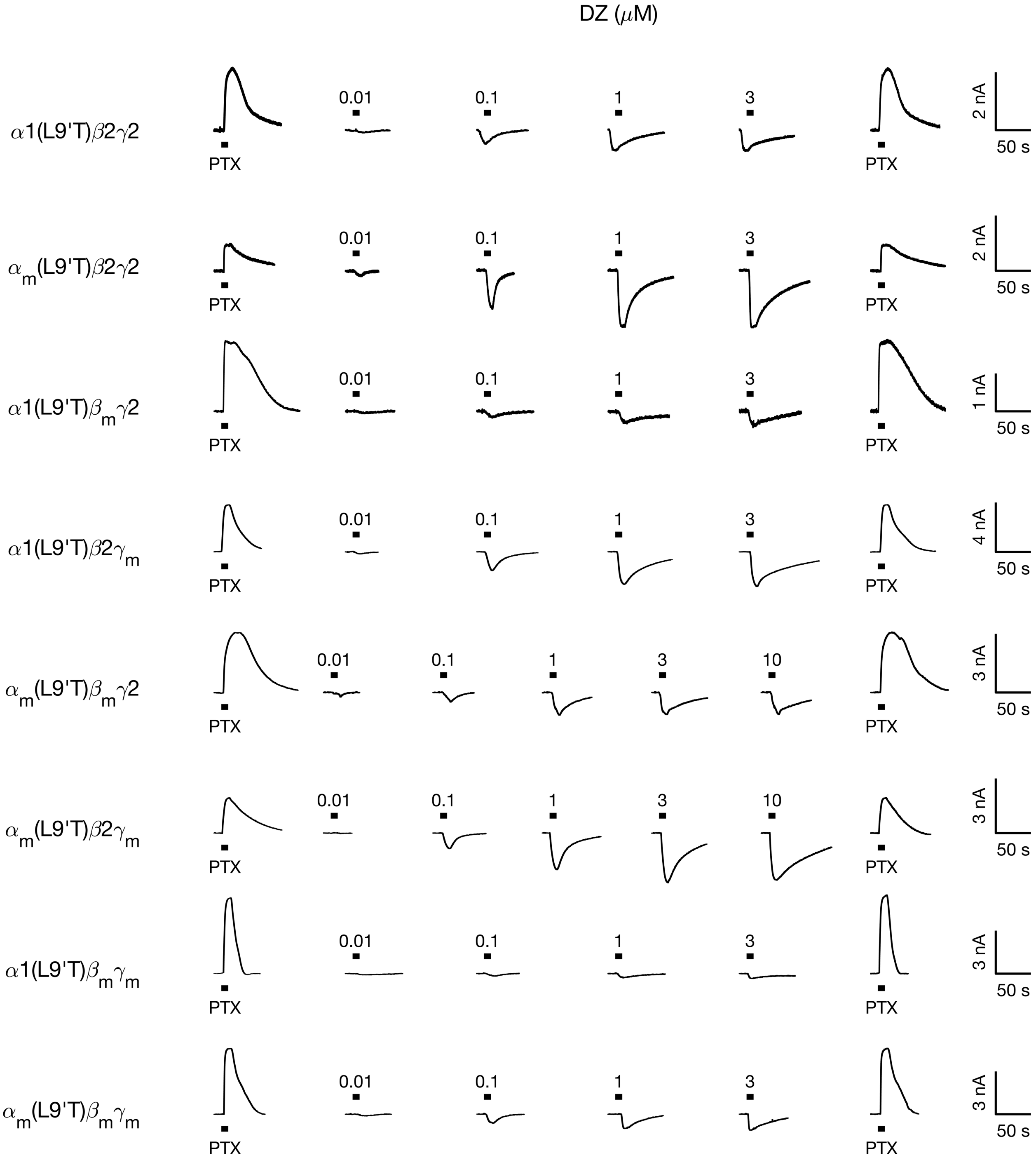


**Figure S2. Example DZ-evoked currents from receptors with M2-M3 linker mutations $\alpha_m$, $\beta_m$, and/or $\gamma_m$ in the gain-of-function α1(L9'T)β2γ2 background.** Currents are in response to 10 second pulses (black bars) of either 1 mM PTX or a series of increasing concentrations of DZ (concentration in micromolar above respective black bar). Summaries of concentration-response curves across oocytes are shown in **Figure S5**. Responses to DZ were bookended by the pore blocker PTX to assess the amount of spontaneous unliganded activity and identify the zero-current baseline.

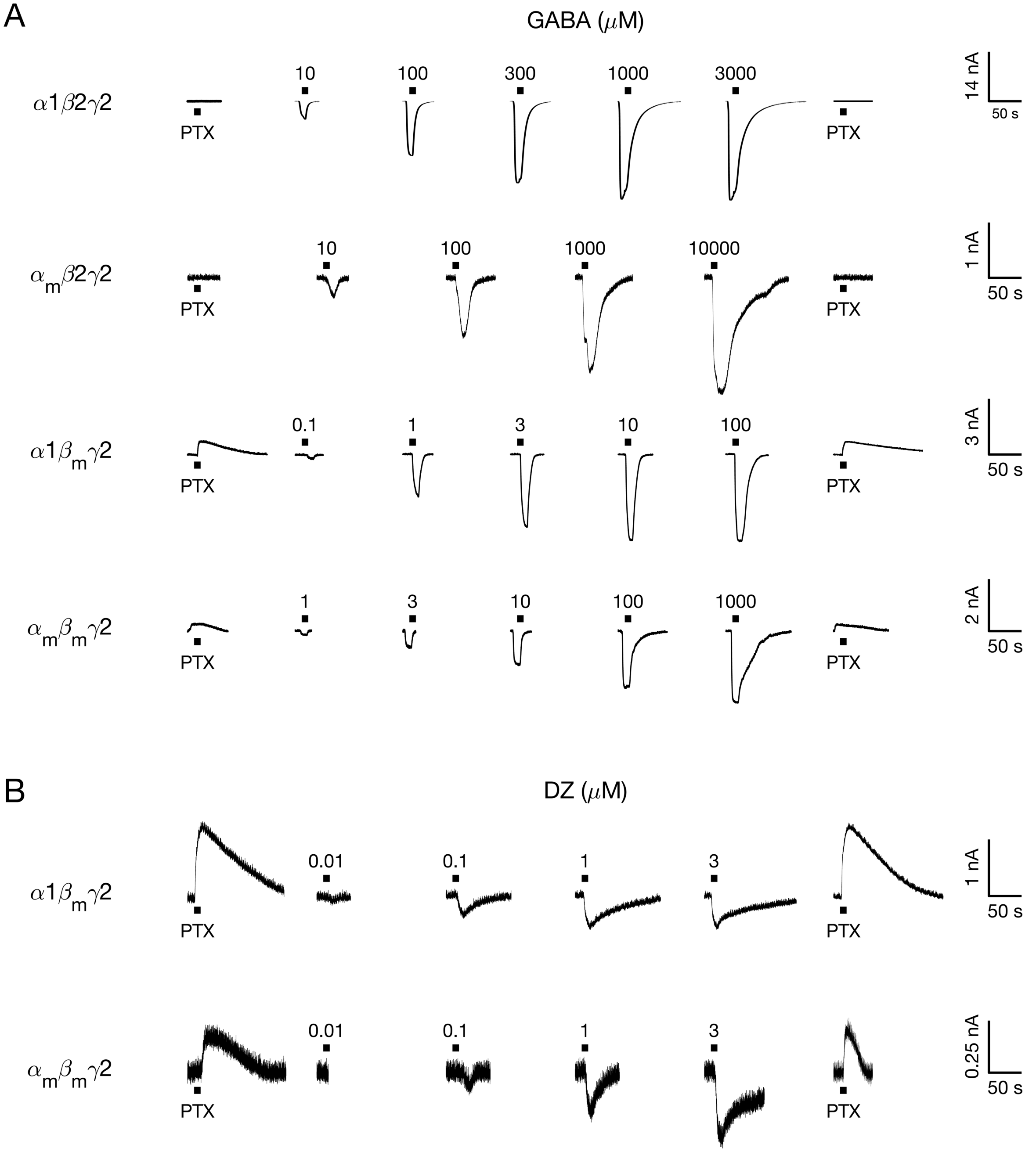


**Figure S3. Example GABA- or DZ-evoked currents from receptors with M2-M3 linker mutations $\alpha_m$, $\beta_m$, and/or $\gamma_m$ in a wild type α1β2γ2 background. (A)** Currents are in response to 10 second pulses (black bars) of either 1 mM PTX or a series of increasing concentrations of GABA (concentration in micromolar above respective black bar). Summaries of concentration-response curves across oocytes are shown in **Figure S6**. Responses to GABA were bookended by the pore blocker PTX to assess the amount of spontaneous unliganded activity and identify the zero-current baseline. **(B)** Same as in **(A)** except for currents evoked with DZ instead of GABA.

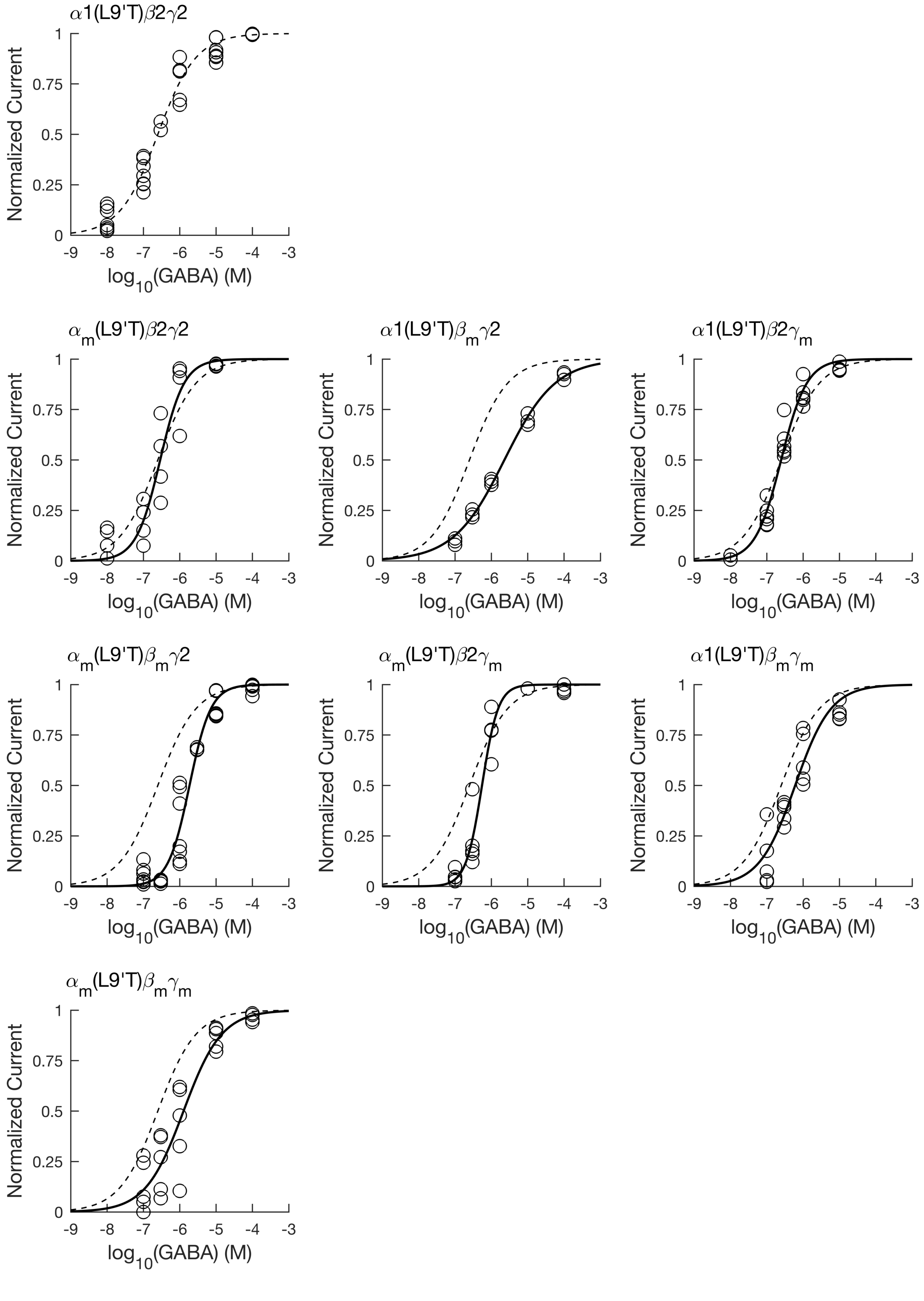


**Figure S4. GABA-evoked concentration-response relationships for gain-of-function α1(L9'T)β2γ2 receptors without and with M2-M3 linker mutations $\alpha_m$, $\beta_m$, and/or $\gamma_m$.** For each oocyte, GABA-evoked current amplitudes from the unliganded baseline as shown in **Figure S1** versus GABA concentration were fit to the Hill equation **(Equation 1)**. Parameters for individual fits are shown in **Figure S7**. Data points for each oocyte where normalized to their Hill fit maximum, and the combined normalized data from all oocytes (circles) was fit to the Hill equation with $I_{max} = 1$ (solid lines, dashed line is fit for $\alpha$1(L9'T)$\beta$2$\gamma$2 receptors). Combined fit parameters ($EC_{50}$, Hill coefficient *h*) and number of oocytes (n) for each construct: $\alpha$1(L9'T)$\beta$2$\gamma$2: $EC_{50}$ = 0.25 $\mu$M, *h* = 0.83, n = 7, $\alpha_m$(L9'T)$\beta$2$\gamma$2: $EC_{50}$ = 0.29 $\mu$M, *h* = 1.31, n = 4, $\alpha$1(L9'T)$\beta_m\gamma$2: $EC_{50}$ = 2.28 $\mu$M, *h* = 0.62, n = 3, $\alpha$1(L9'T)$\beta$2$\gamma_m$: $EC_{50}$ = 0.25 $\mu$M, *h* = 1.23, n = 6, $\alpha_m$(L9'T)$\beta_m\gamma$2: $EC_{50}$ = 1.86 $\mu$M, *h* = 1.47, n = 7, $\alpha_m$(L9'T)$\beta$2$\gamma_m$: $EC_{50}$ = 0.53 $\mu$M, *h* = 1.95, n = 5, $\alpha$1(L9'T)$\beta_m\gamma_m$: $EC_{50}$ = 0.6 $\mu$M, *h* = 0.88, n = 5, $\alpha_m$(L9'T)$\beta_m\gamma_m$: $EC_{50}$ = 1.27 $\mu$M, *h* = 0.83 , n = 5.

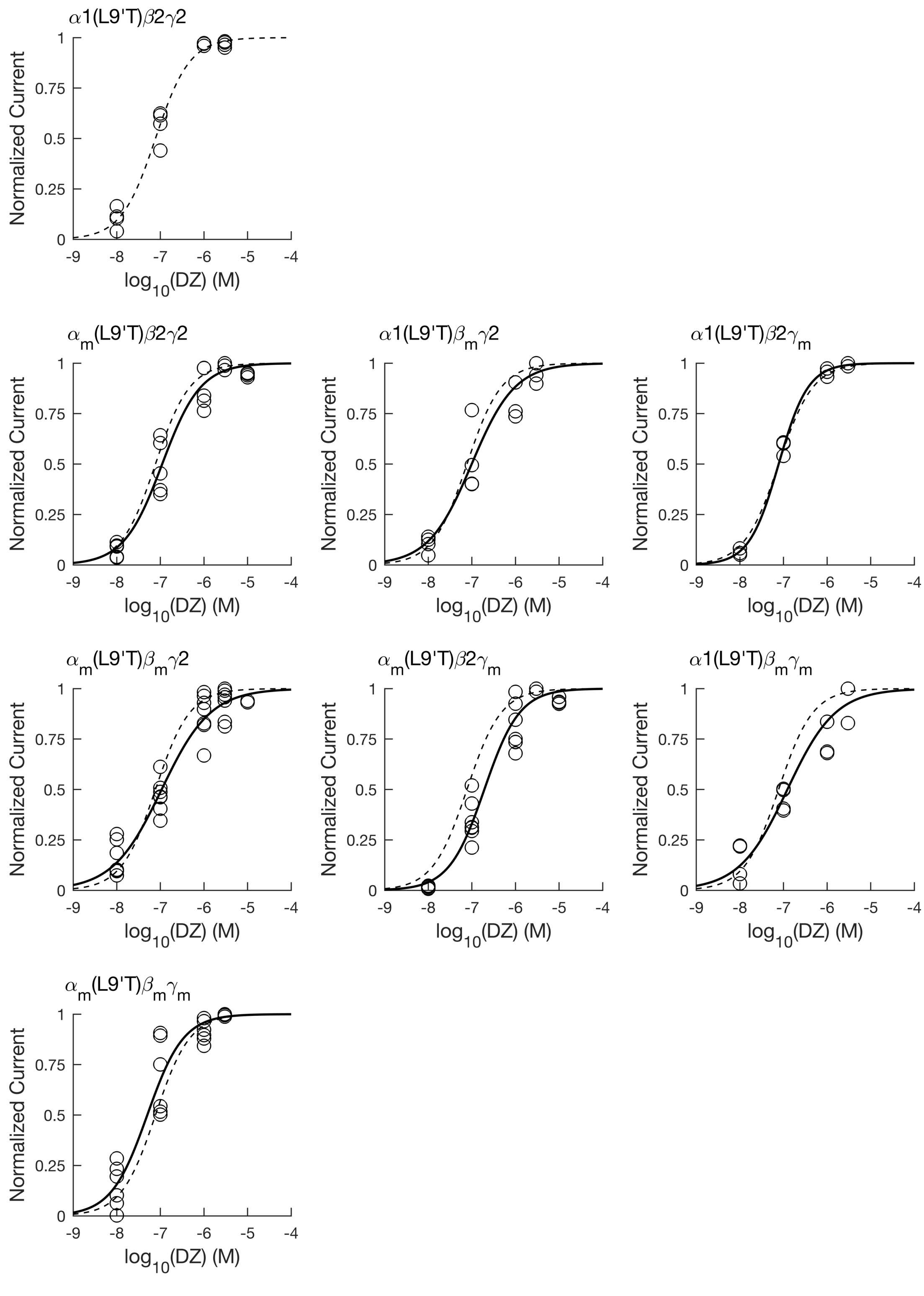


**Figure S5. DZ-evoked concentration-response relationships for gain-of-function $\alpha$1(L9'T)$\beta$2$\gamma$2 receptors without and with M2-M3 linker mutations $\alpha_m$, $\beta_m$, and/or $\gamma_m$.** For each oocyte, DZ-evoked current amplitudes from the unliganded baseline as shown in **Figure S2** versus DZ concentration were fit to the Hill equation **(Equation 1)**. Parameters for individual fits are shown in **Figure S8**. Data points for each oocyte where normalized to their Hill fit maximum, and the combined normalized data from all oocytes (circles) was fit to the Hill equation with $I_{max} = 1$ (solid lines, dashed line is fit for $\alpha$1(L9'T)$\beta$2$\gamma$2 receptors). Combined fit parameters ($EC_{50}$, Hill coefficient *h*) and number of oocytes (n) for each construct: $\alpha$1(L9'T)$\beta$2$\gamma$2: $EC_{50}$ = 0.08 $\mu$M, *h* = 1.10, n = 4, $\alpha_m$(L9'T)$\beta$2$\gamma$2: $EC_{50}$ = 0.11 $\mu$M, *h* = 0.98, n = 5, $\alpha$1(L9'T)$\beta_m\gamma$2: $EC_{50}$ = 0.1 $\mu$M, *h* = 0.88, n = 4, $\alpha$1(L9'T)$\beta$2$\gamma_m$: $EC_{50}$ = 0.08 $\mu$M, *h* = 1.28, n = 3, $\alpha_m$(L9'T)$\beta_m\gamma$2: $EC_{50}$ = 0.11 $\mu$M, *h* = 0.77, n = 7, $\alpha_m$(L9'T)$\beta$2$\gamma_m$: $EC_{50}$ = 0.19 $\mu$M, *h* = 1.07, n = 7, $\alpha$1(L9'T)$\beta_m\gamma_m$: $EC_{50}$ = 0.12 $\mu$M, *h* = 0.79, n = 4, $\alpha_m$(L9'T)$\beta_m\gamma_m$: $EC_{50}$ = 0.05 $\mu$M, *h* = 1.05, n = 6.

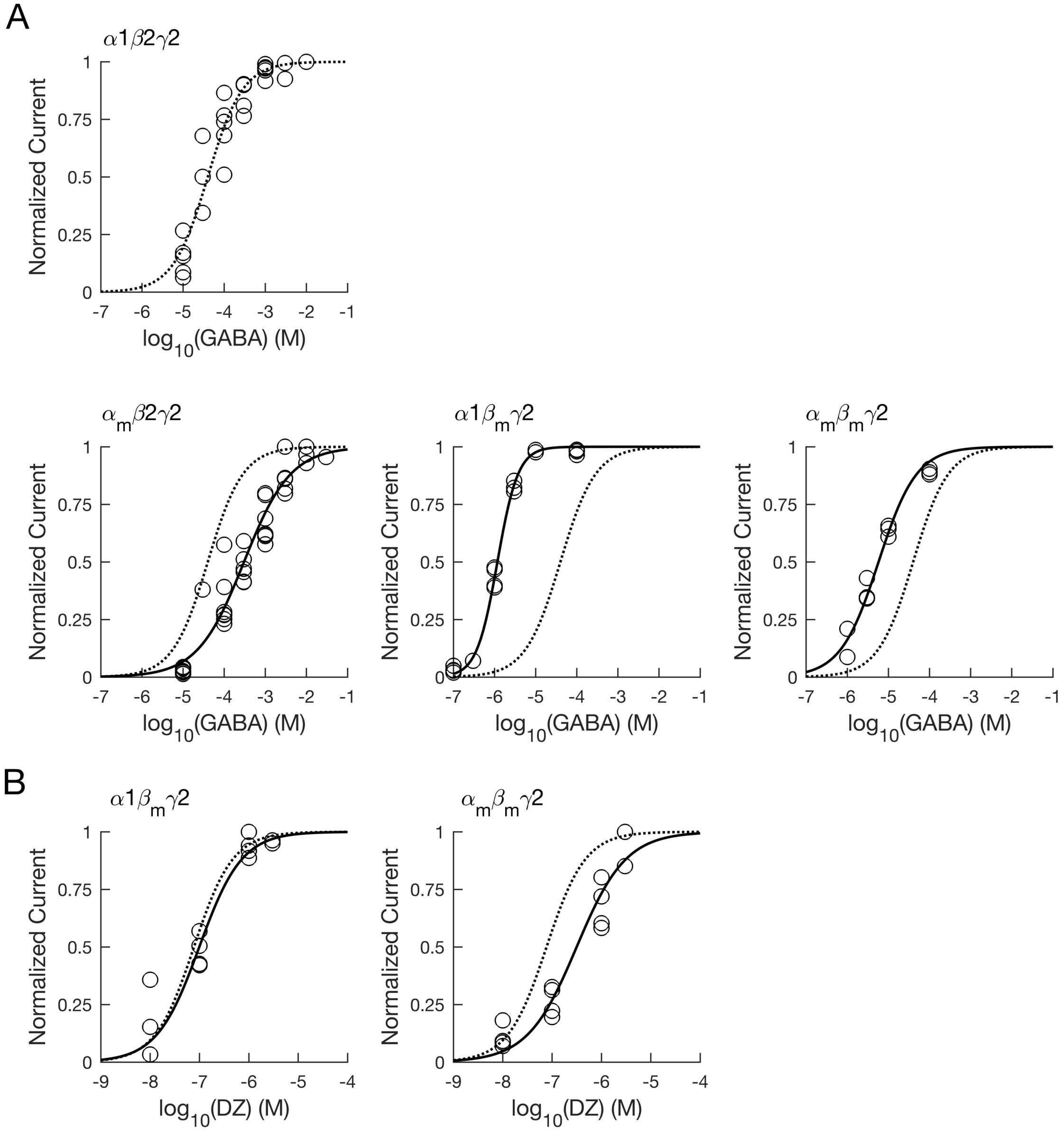


**Figure S6. GABA- and DZ-evoked concentration-response relationships for wild type α1β2γ2 receptors without and with M2-M3 linker mutations $\alpha_m$, $\beta_m$, and/or $\gamma_m$.** For each oocyte, GABA- **(A)** or DZ-evoked **(B)** current amplitudes from the unliganded baseline as shown in **Figure S3** versus GABA or DZ concentration, respectively, were fit to the Hill equation **(Equation 1)**. Parameters for individual fits are shown in **Figures S7-S8**. Data points for each oocyte where normalized to their Hill fit maximum, and the combined normalized data from all oocytes (circles) was fit to the Hill equation with $I_{max} = 1$ (solid lines, dashed line is fit for α1β2γ2 receptors). GABA-evoked **(A)** combined fit parameters ($EC_{50}$, Hill coefficient *h*) and number of oocytes (n) for each construct: α1β2γ2: $EC_{50}$ = 39.94 μM, *h* = 1.02, n = 5, $\alpha_m\beta 2\gamma 2$: $EC_{50}$ = 303.37 μM, *h* = 0.77, n = 7, $\alpha 1\beta_m\gamma 2$: $EC_{50}$ = 1.17 μM, *h* = 1.73, n = 4, $\alpha_m\beta_m\gamma 2$: $EC_{50}$ = 5.76 μM, *h* = 0.94, n = 3. DZ-evoked **(B)** combined fit parameters ($EC_{50}$, Hill coefficient *h*) and number of oocytes (n) for each construct: $\alpha 1\beta_m\gamma 2$: $EC_{50}$ = 0.1 μM, *h* = 1.01, n = 4, $\alpha_m\beta_m\gamma 2$: $EC_{50}$ = 0.31 μM, *h* = 0.87, n = 4.

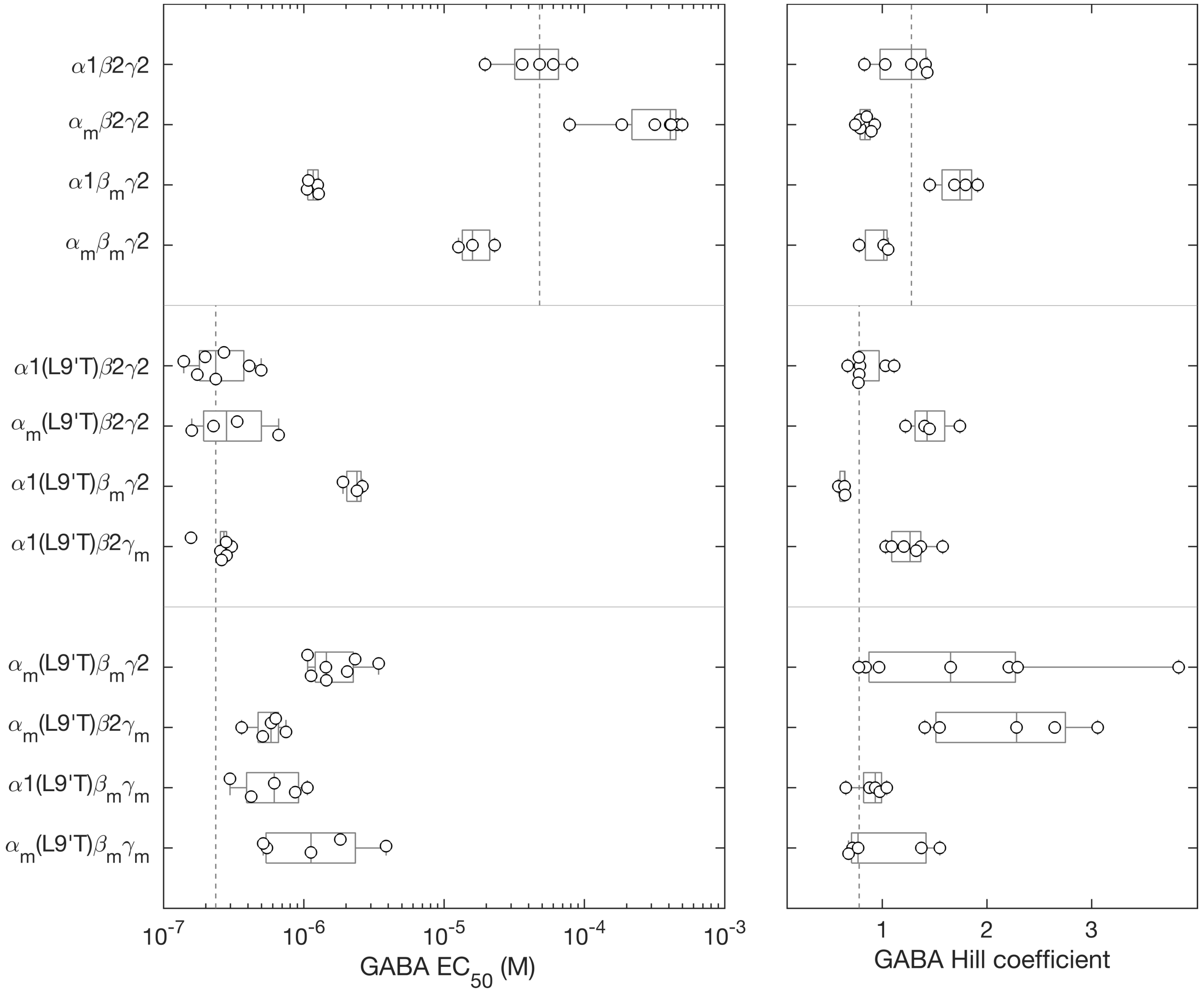


**Figure S7. Summary of GABA concentration-response relation Hill fit parameters for individual oocytes.** Data points are for individual oocytes ($\alpha1\beta2\gamma2$: n = 5; $\alpha_m\beta2\gamma2$: n = 7; $\alpha1\beta_m\gamma2$: n = 4; $\alpha_m\beta_m\gamma2$: n = 3; $\alpha1$(L9'T)$\beta2\gamma2$: n = 7; $\alpha_m$(L9'T)$\beta2\gamma2$: n = 4 ; $\alpha1$(L9'T)$\beta_m\gamma2$: n = 3; $\alpha1$(L9'T)$\beta2\gamma_m$: n = 6; $\alpha_m$(L9'T)$\beta_m\gamma2$: n = 7 ; $\alpha_m$(L9'T)$\beta2\gamma_m$: n = 5 ; $\alpha1$(L9'T)$\beta_m\gamma_m$: n = 5 ; $\alpha_m$(L9'T)$\beta_m\gamma_m$: n = 5). Box plots indicate quartiles, and the upper and lower vertical dashed lines are the median for the $\alpha1$(L9'T)$\beta2\gamma2$ and $\alpha1\beta2\gamma2$ backgrounds, respectively.

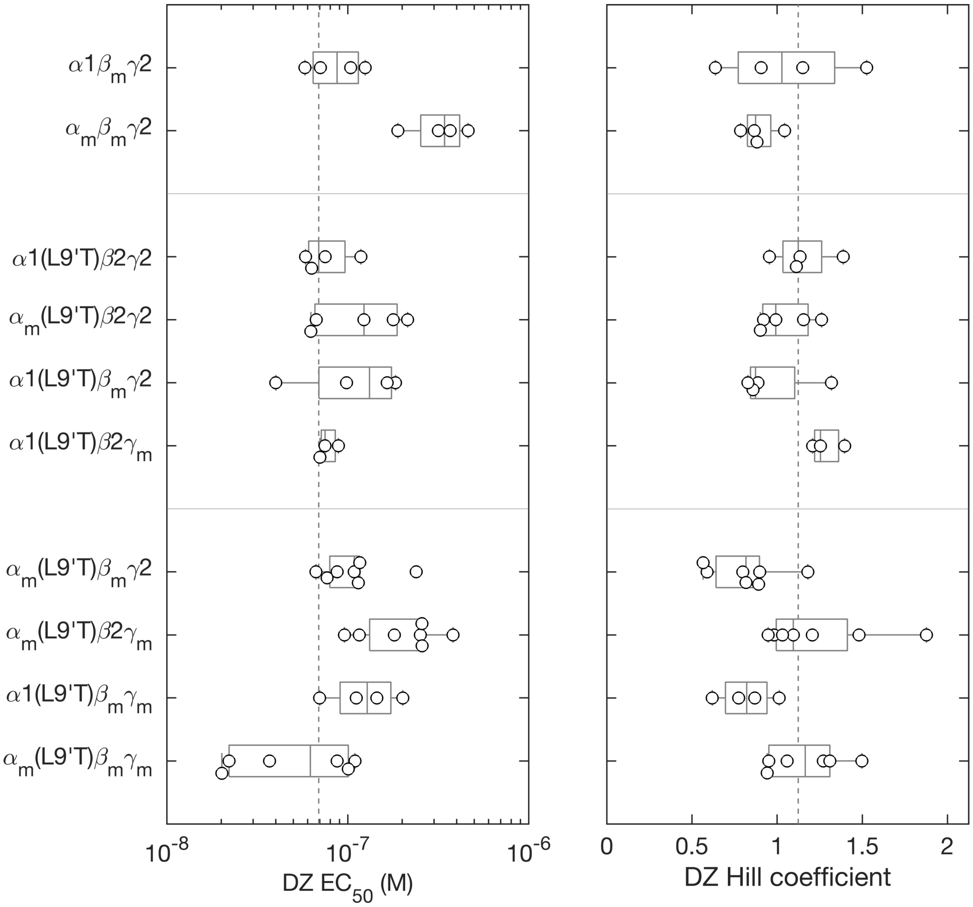


**Figure S8. Summary of DZ concentration-response relation Hill fit parameters for individual oocytes.** Data points are for individual oocytes ($\alpha1\beta_m\gamma2$: n = 4; $\alpha_m\beta_m\gamma2$: n = 4; $\alpha1$(L9'T)$\beta2\gamma2$: n = 4; $\alpha_m$(L9'T)$\beta2\gamma2$: n = 5; $\alpha1$(L9'T)$\beta_m\gamma2$: n = 4; $\alpha1$(L9'T)$\beta2\gamma_m$: n = 3; $\alpha_m$(L9'T)$\beta_m\gamma2$: n = 7; $\alpha_m$(L9'T)$\beta2\gamma_m$: n = 7; $\alpha1$(L9'T)$\beta_m\gamma_m$: n = 4; $\alpha_m$(L9'T)$\beta_m\gamma_m$: n = 6). Box plots indicate quartiles, and the vertical dashed line is the median for the $\alpha1$(L9'T)$\beta2\gamma2$ background.

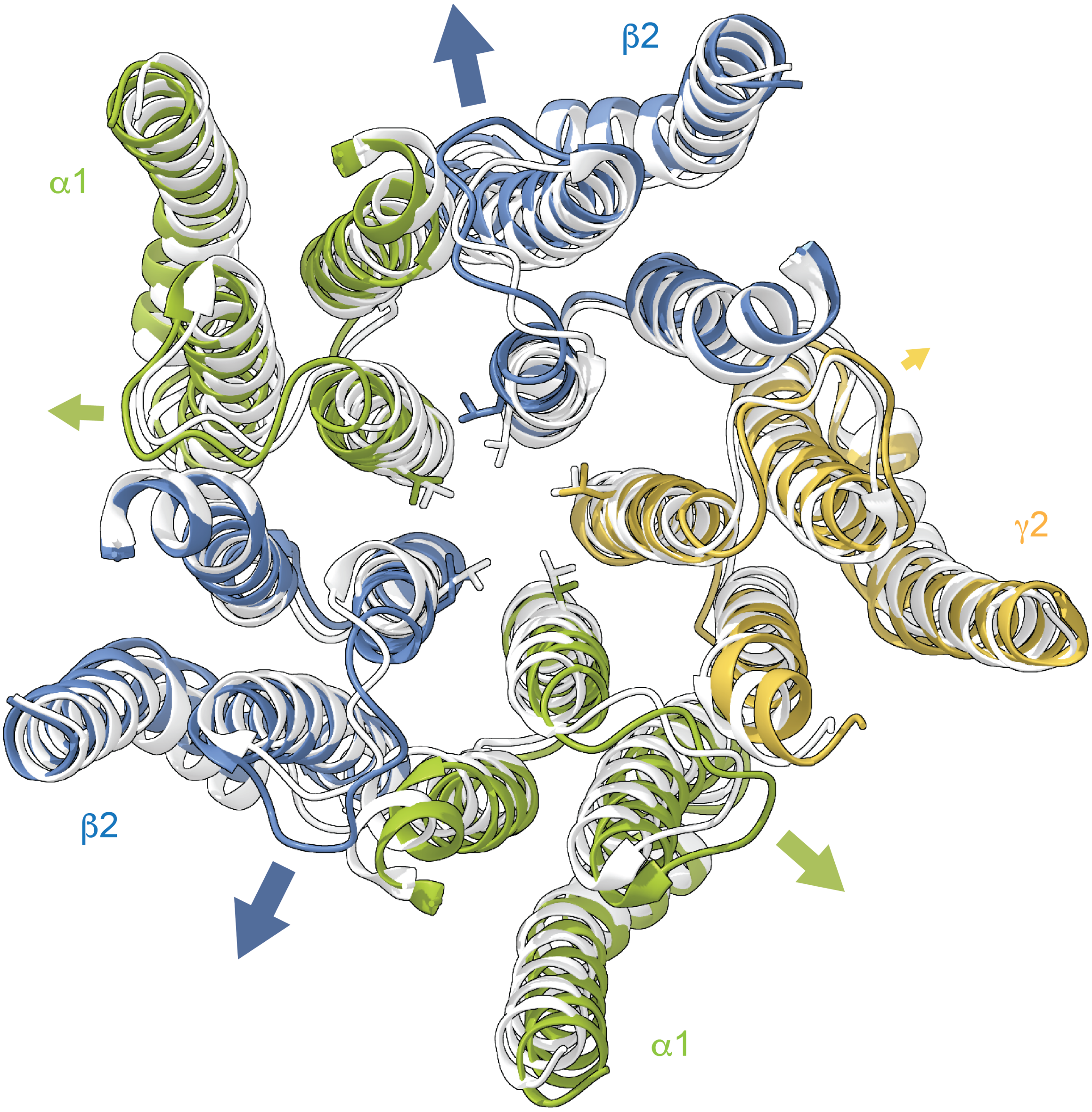


**Figure S9. Asymmetric M2-M3 linker and M2 helix motions during channel gating.** Comparison of the TMD for cryo-EM structures of α1β2γ2 receptors in complex with either the antagonist bicuculline (light gray, closed conformation; PDB 6X3S) or GABA (colored subunits, desensitized conformation; PDB 6X3Z). TMD helices were aligned in ChimeraX (Pettersen et al., 2021). The ECD is omitted for clarity. Top-down view looking through the channel into the cell. The 9' leucines forming the central pore gate are shown as sticks. Comparison of closed versus desensitized conformations indicative of motions occurring during pore opening indicates a relatively larger radial displacement of the M2-M3 linker and rotation of the 9' leucine out of the conducting pathway in β2 subunits as compared to α1 or γ2 subunits (arrows).
